# Single-cell chromatin landscapes visualize epigenetic barriers and reveal lineage-specific Polycomb-mediated repression

**DOI:** 10.64898/2026.05.04.722686

**Authors:** Sergei Pirogov, Aleksander Purik, Artem Ilin, Jose Ramon Barcenas Walls, Marek Bartosovic, Mattias Mannervik

## Abstract

Understanding how chromatin state contributes to developmental trajectories remains central to deciphering cell specification and differentiation. Using dual-modality nano-CUT&Tag, we profiled two antagonistic histone modifications—active H3K27ac and repressive H3K27me3—in thousands of single cells from Drosophila embryos across early lineage diversification and terminal differentiation. Joint embedding of both marks enabled robust cell-type classification and revealed increasing epigenetic specificity over developmental time. We ordered cells by developmental age and epigenomic similarity, and defined an epigenetic potential metric that visualizes repressive chromatin barriers as landscapes that predict transcriptional activity. While many genes conform to a classical model in which expression resides in low-potential epigenetic valleys, a substantial subset shows co-occurrence of H3K27ac, H3K27me3, and transcription within the same cell lineage. This indicates that Polycomb-mediated H3K27me3 repression frequently acts within, rather than solely between, lineages. Consistently, tissue-specific E(z) knockdown demonstrates that partial loss of H3K27me3 predominantly de-represses lineage-matched genes rather than inducing fate conversion. Systematic analysis showed that H3K27me3 occurs in multiple distributional modes, ranging from ubiquitous to highly cell-type-specific deposition, co-occuring with accessible but silent gene promoters. These findings demonstrate that cell-type-specific deployment of H3K27ac and H3K27me3 sculpts epigenetic potential landscapes that shape developmental gene expression patterns.

## Introduction

Understanding how cells progressively acquire specific identities and differentiate is a fundamental goal of developmental biology. During this process, cells integrate intrinsic regulatory programs with context-specific cues such as niche interactions, intercellular signaling, and spatial position (Morris, 2019; Rafelski and Theriot, 2024). Cell specification and differentiation are guided by gene regulatory networks (GRNs) that are often arranged hierarchically (Levine and Davidson, 2005). At the top of these hierarchies are “master regulatory” transcription factors that initiate regulatory cascades by activating secondary transcription across different tissues (Balsalobre and Drouin, 2022; Larson et al., 2021; Spitz and Furlong, 2012). Transcription factors further facilitate recruitment of co-activators and co-repressors, which create chromatin states that are either permissive or restrictive to gene expression (Mannervik et al., 1999). Waddington’s epigenetic landscape provides a useful conceptual framework for this process, depicting developmental trajectories as valleys surrounded by hills that help stabilize cell identity (Waddington, 1957).

Two key opposing histone modifications are associated with gene regulatory functions. Histone 3 lysine 27 acetylation (H3K27ac) is found at active genes and cis-regulatory elements (Creyghton et al., 2010; Rada-Iglesias et al., 2011), whereas H3K27 tri-methylation (H3K27me3) is associated with facultative gene repression during development (Trojer and Reinberg, 2007). In Drosophila, H3K27ac is deposited by the histone acetyltransferase CBP (also known as Nejire), the sole orthologue of mammalian co-activators p300 and CBP, whereas H3K27me3 is deposited by Enhancer of zeste, E(z), the EZH1/2 orthologue within Polycomb Repressive Complex 2, PRC2 (Bannister and Kouzarides, 1996; Cao et al., 2002; Czermin et al., 2002; Tie et al., 2009). Polycomb-mediated H3K27 methylation establishes repressive barriers that contribute to epigenetic homeostasis (Adam and Fuchs, 2016; Flavahan et al., 2017). When these barriers are weakened, lineage fidelity erodes, potentially leading to disease states, including tumorigenesis (Feinberg and Levchenko, 2023; Flavahan et al., 2017; Gronkowska and Robaszkiewicz, 2024; Parreno et al., 2024).

Nevertheless, the emergence of chromatin barriers between cell lineages during Drosophila embryonic development remains poorly understood. To investigate this process, we profiled H3K27ac and H3K27me3 in thousands of single cells from Drosophila embryos. We utilized nano-CUT&Tag to simultaneously map both marks with single cell resolution at two critical stages of Drosophila embryogenesis: during cell specification/diversification and during terminal differentiation. We then developed a novel computational approach for chromatin state visualization inspired by Waddington’s epigenetic landscape. These landscapes illustrate differences in epigenetic potential between developing cell lineages, providing a framework to examine how the dynamics of H3K27ac and H3K27me3 restrict gene expression and cell fate. We found that many loci carry H3K27me3 in tissues where they are inactive, as expected. In addition, a substantial set of genes showed cell-type-restricted H3K27me3 in the lineage where the gene is normally active. This pattern suggests that Polycomb-mediated repression creates barriers between closely related cell types, where regulatory crosstalk would otherwise be likely. Consistent with this idea, partial depletion of H3K27me3 de-represses Polycomb-target genes within a lineage, indicating that these loci are primed to respond to activation cues once the repressive barrier is lowered.

## Results

### Dual epigenome profiling achieves cell-type resolution in Drosophila embryos

To map the co-dynamics of histone marks associated with gene activation and repression across *Drosophila melanogaster* embryogenesis, we employed single-cell nano-CUT&Tag (Barcenas-Walls et al., 2024; Bartosovic and Castelo-Branco, 2023). This method enables simultaneous mapping of two chromatin marks using barcoded nanobody-Tn5 transposase recognizing primary antibodies raised in different species (Fig. 1A), allowing for simultaneous profiling of both marks in the same cells.

**Figure 1.**
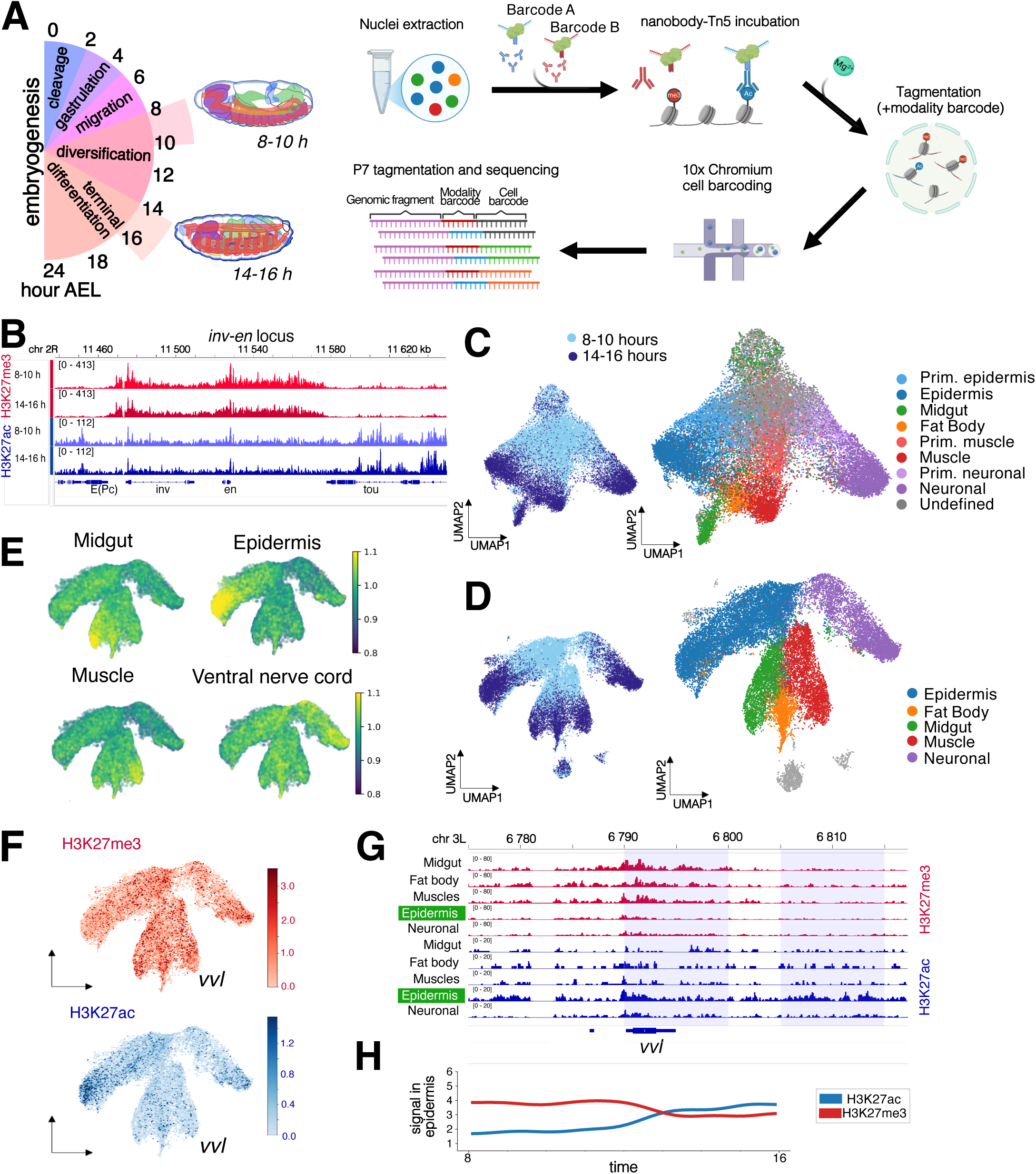
nanoCUT&Tag resolves cell type-specific chromatin states in the Drosophila embryo. **A.** Experimental workflow. Drosophila embryos were collected at two developmental windows, 8–10 h and 14–16 h after egg laying (AEL), dissociated to nuclei, and incubated with nanobody–Tn5 complexes targeting H3K27ac and H3K27me3. After tagmentation and 10x Genomics barcoding, libraries were sequenced and demultiplexed into a bimodal cell×genomic bin matrix. **B.** Genome browser snapshots showing pseudobulk H3K27ac (blue) and H3K27me3 (red) at the *inv–en* locus for 8–10 h and 14–16 h embryos. **C.** UMAP visualizations of nanoCUT&Tag nuclei from two biological replicates. Left, two-mark cell×bin embedding colored by collection windows. Right, the same embedding colored by Leiden clusters annotated to major tissues by anatomic and GO-term enrichment for top cluster-specifically acetylated genes. **D.** UMAP visualization of nanoCUT&Tag nuclei after integration with published 8–16 h scATAC-seq (Calderon et al., 2022). Left, integrated UMAP colored by collection windows. Right, integrated UMAP colored by annotated clusters; cell clusters not considered for the downstream analysis are shown in light grey. **E.** Tissue gene-set projections on the integrated UMAP. For each panel (midgut, epidermis, muscle, ventral nerve cord), per-cell H3K27ac signal was summed over bins linked to genes annotated with the corresponding anatomical term (n = epidermis: 623, muscle: 608, midgut: 999, vnc: 896 genes). **F.** Per-cell projection of signals for the epidermal transcription factor *vvl*, H3K27me3 (red) and H3K27ac (blue) signals summed over genomic bins linked to *vvl*. **G.** Pseudobulk genome browser snapshot of H3K27ac/H3K27me3 at *vvl* across the main cell types at 14–16 h. The connected bins are shown with light grey background. The cell type with major gene expression is highlighted with a green box. **H.** H3K27ac and H3K27me3 signal change at *vvl* during inferred developmental time (8-16 h AEL) in the epidermal cluster.

We applied nano-CUT&Tag at two distinct developmental time points, 8-10 hours after egg laying (AEL, stages 12-13), during cell specification/diversification, and 14-16 hours AEL (stage 16), representing terminal differentiation. After quality control, we retrieved 27,599 cells from the two stages and two biological replicates that correlated well with each other (Fig. S1A-C, Table S1). We obtained 112-265 mean unique fragments per cell for H3K27me3, and 211-467 fragments per cell for H3K27ac (Fig. S1D, E), and pseudobulk analysis showed expected patterns with H3K27me3 forming broad domains, while H3K27ac was enriched on gene promoters and enhancers (Fig. 1B).

To co-embed both modalities, we used the multi-view spectral embedding provided by the SnapATAC2 package (Zhang et al., 2023). To avoid any loss of information, we used a cell x 5kb-long bin matrix of the genome, which performed better than a cell x peak matrix (Fig. S1F). Co-embedded data formed developmental trajectories, with the earlier stage enriched in the trunk and later-stage cells forming diverging tips of the branches (Fig. 1C). We observed that the late developmental stage (14-16 h AEL) showed better separation of cell types in 2D Uniform Manifold Approximation and Projection (UMAP)-transformed space than the earlier stage 8-10 h AEL (Fig. S2A). The multi-view spectral embedding of both histone marks outperformed unimodal embeddings for H3K27ac and H3K27me3, highlighting the additional sensitivity provided by multimodal profiling. Nevertheless, 14-16 h AEL cells could be resolved on the UMAP based on the individual histone modalities (Fig. S2B).

In order to annotate the cell clusters, we used gene ontology (GO)-terms enriched in genes with cluster-specific H3K27 acetylation, in combination with developmental anatomy terms extracted from the BDGP in situ database (Tomancak et al., 2002). We used a regulatory inference graph of gene-bin connections to link between one and ten bins per gene (see Methods and Fig. S2C, D), and identified epidermal, muscle, fat body, midgut, and neuronal cell clusters, as well as primordia of epidermal, muscle, and neuronal lineages (Fig.1C, S2A, B, Table S2). We were not able to annotate one cell cluster, which may consist of yolk, amnioserosa, and undifferentiated nuclei. To enhance cluster resolution, we integrated our nano-CUT&Tag data with a published scATAC-seq atlas that covers the entire course of embryogenesis (Calderon et al., 2022), using scGLUE (Cao and Gao, 2022). The scATAC-seq data served as an effective scaffold, improving trajectory visualization and clustering, and supporting cluster annotation (Fig. 1D, S3A,B). The consistency of clusters before and after integration with scATAC shows that cluster-specific overlap between H3K27ac and H3K27me3 is not an artifact of integration (Fig. S3B). We confirmed cell-type specificity by projecting H3K27ac signal on the UMAP for gene-sets enriched in each cell cluster (Fig. 1E). Known cell-type-specific genes, such as the transcription factors *ventral veins lacking* (*vvl*) and *grainy head* (*grh*) (expressed in epidermis), *Oli* (neuronal), *Myocyte enhancer factor 2* (*Mef*2) (expressed in muscle), and *GATAe* (endodermal), show enrichment of acetylation in relevant clusters (Fig. 1F, S3C). Pseudobulk genome browser tracks showed that the *vvl* locus is methylated in midgut, fat body and muscles, but most strongly acetylated in epidermis where it is expressed (Fig. 1G). We conclude that the profiles of H3K27ac and H3K27me3 are able to separate major cell types in the Drosophila embryo.

### Developmental dynamics of the epigenome

We next explored the developmental dynamics of these histone marks between the two embryo stages, observing genes with cluster-specific loss and gain of H3K27ac at their respective bins (Fig. S4A, C). For example, in epidermis we identified 1686 genes that lost H3K27ac and 1790 genes that gained acetylation during late embryogenesis (Fig. S4C). For H3K27me3, we observed preferential gain of the mark at the later stage, with an enrichment of transcription factor genes (Fig. S4B, D, Table S3). Four hundred thirty four genes changed both H3K27ac and H3K27me3 status from 8-10h to 14-16h embryos (Fig. S4E).

An increased cell-type specificity was observed for H3K27ac during development, as seen in stronger cluster-specific H3K27 acetylation at the late stage for the top 200 genes used for cluster annotation (Fig. S5A), illustrated in genome browser shots for the *vvl*, *oli* and *srp-GATAe* loci (Fig. S5B). We quantified this increased specificity by comparing Shannon entropies between early and late stages (Fig. S5C, D). Consistent with this, we found that the correlation of H3K27-acetylated or -methylated genes between cell types decreased during development, reflecting divergence of epigenomes during tissue differentiation (Fig. S5E). The neuronal cell lineage was the most divergent, both in terms of H3K27 acetylation and methylation.

We next visualized the overall dynamics of H3K27ac and H3K27me3 for specific genes (*GATAe, Mef2, vvl, grh* and *Oli)* by integration with developmental time from the scATAC-seq dataset (Fig.1H, S5F), illustrating that cell-type specific gene activity is accompanied by an increase in H3K27me3 and by higher specificity of H3K27ac during differentiation. This explains the better cell type separation at the late stage in the UMAP space. Taken together, the data demonstrates increased epigenetic specificity over developmental time.

### A cell type-developmental time embedding for chromatin landscapes

To facilitate assessment of histone mark distribution and dynamics, we designed a novel cell type-developmental time embedding (Fig. 2A). This approach arranges cells along one axis by their epigenomic similarity and along the other by their inferred developmental time, facilitating the side-by-side comparison of cell trajectories. We created a scaffold by making an ordered dendrogram of late embryonic stage cell clusters (Fig. 2A, S6A). Inferred developmental time was assigned to the cells through integration with time-resolved scATAC-seq data, where each cell received a neural network-predicted age (Fig. 2B) (Calderon et al., 2022). The inferred developmental time aligns well with our embryo collection time windows, despite the presence of cells with an estimated age between the collection windows. Then, we grouped all cells into metacells and embedded them according to their similarity to each dendrogram tip, and to their average inferred developmental time (Fig. S6B). The individual cells were distributed along the x-axis depending on the position of the metacell, and on the y-axis according to inferred cell age. Cells were then allowed to find their final positions by running a limited number of UMAP iterations (Fig. S6C), and labeled according to cell type (Fig. 2B).

**Figure 2.**
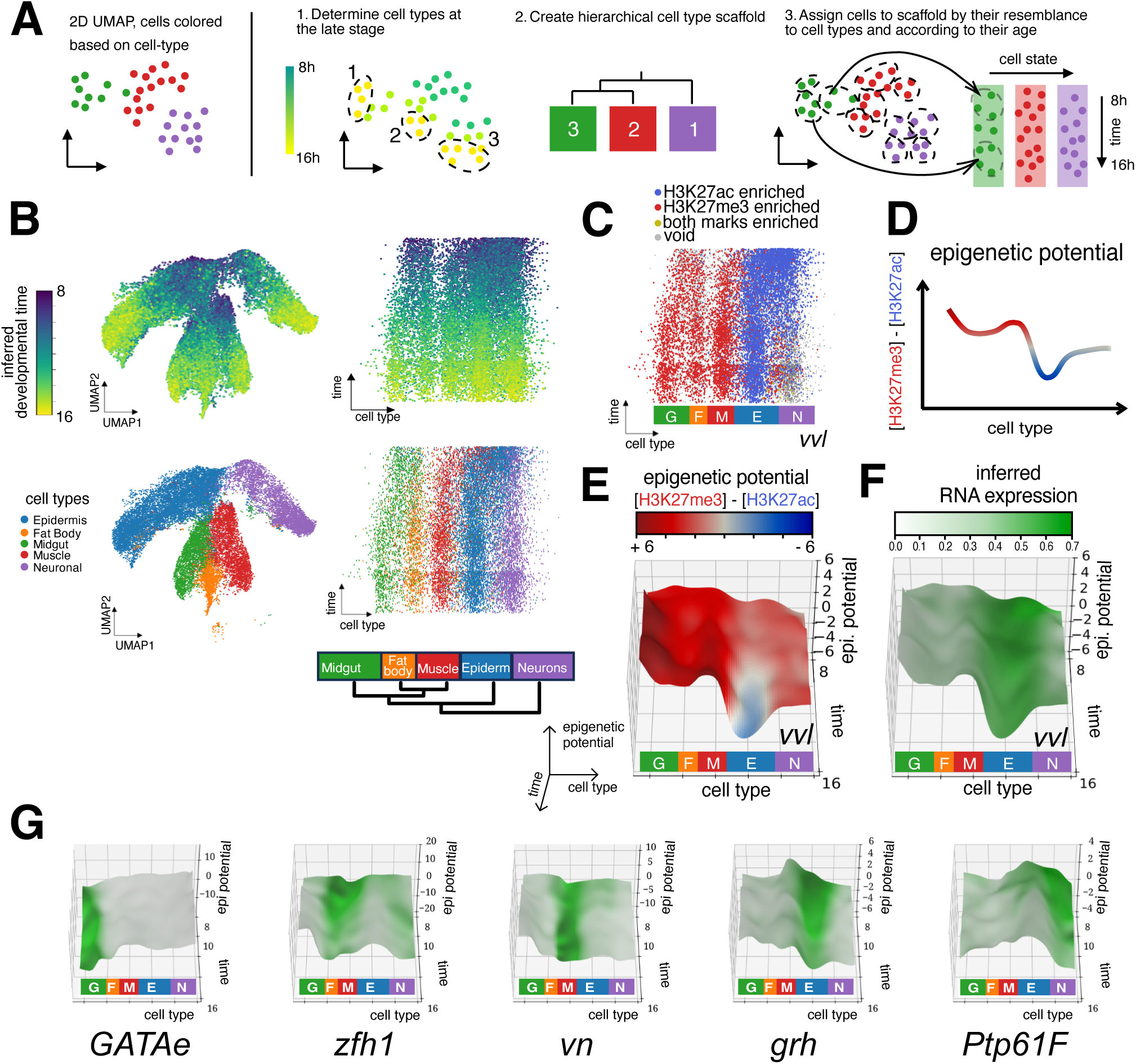
Gene-specific epigenetic landscapes visualize transcriptional permissiveness across cell types and developmental time. **A.** Schematic of landscape construction. **B.** Top left, developmental time transferred from the integrated scATAC-seq (Calderon et al., 2022) projected onto the nanoCUT&Tag UMAP. Bottom left, the same UMAP colored by cell-type annotation (as in Fig.1D). Top right, inferred developmental time projected onto the cell type-developmental time embedding. Bottom right, cell type–developmental time embedding colored by cell type (Epidermis, Fat body, Midgut, Muscle, Neuronal). The dendrogram shows similarity between cell types. **C.** Cell type–developmental time embedding colored by classification of chromatin signal at bins linked to *vvl*: H3K27ac-enriched, H3K27me3-enriched, both marks enriched, or void. Midgut (G), Fat body (F), Muscle (M), Epidermis (E), Neuronal (N). **D.** Epigenetic potential diagram, showing one time slice of an epigenetic landscape, where epigenetic potential is calculated as H3K27ac signal subtracted from H3K27me3 signal. **E.** Epigenetic landscape for *vvl.* Surface plot over the cell–time plane where height (z-axis) is epigenetic potential. Surface is colored by epigenetic potential. **F.** The same surface plot colored by *vvl* expression levels transferred from inferred scRNA-seq (Calderon et al., 2022). High epigenetic potential denotes repressive “hills”, whereas low/negative epigenetic potential forms permissive “valleys” for expression. **G.** 3D epigenetic landscapes for selected genes colored by level of RNA expression.

We next plotted the histone modifications for each gene on this cell type-time embedding, shown for *vvl* in Fig. 2C. In order to classify a gene as being H3K27 acetylated, methylated, having both marks, or none of them in an individual cell, we calculated a community score per cell. This helped to exclude cells with a high signal that are surrounded by cells without a signal (i.e., false positives), and include cells lacking a signal, but surrounded by cells that are heavily modified at that gene (i.e., false negatives) (see Methods and examples in Fig. S6D). The *vvl* gene shows how acetylation and methylation are distributed on the cell-time embedding before (Fig. S6E) and after histone mark enrichment classification (Fig. 2C). This illustrates the distribution of active and repressive marks among cell types.

To track histone mark dynamics over cell types and developmental time, we created chromatin landscapes for individual genes based on their epigenetic potential (hypothetical permissiveness to gene expression). This was based on H3K27me3 (restrictive) minus H3K27ac (permissive) signal in the genomic region allocated to each gene (Fig. 2D). This epigenetic potential is based on signal with full dynamic range, and not on the chromatin mark classification used in Fig. 2C. For the epidermal transcription factor gene *vvl*, the landscape clearly shows a low-potential valley along the epidermal trajectory, reflecting high H3K27ac and low H3K27me3 enrichment (Fig. 2E). Crucially, co-plotted scRNA-seq data from (Calderon et al., 2022) confirmed that high RNA expression follows this low-potential epigenetic valley (Fig. 2F). This demonstrates that our chromatin landscape can be a strong predictor of gene expression, and such landscapes are present also at other genes with cell-type-specific expression (Fig. 2G, S6F). Thus, these chromatin landscapes provide a visualisation framework to illustrate how the co-dynamics of H3K27ac and H3K27me3 may channel gene expression and cell fate during development.

### H3K27ac, RNA expression and H3K27me3 frequently co-occur in the same cell lineage

We observed that whereas expression of many genes negatively correlates with epigenetic potential, as expected, a distinct subset of genes exhibited a positive correlation between high epigenetic potential and gene expression, implying that the presence of repressive marks can co-occur with gene expression (Fig. 3A, B). This co-occurrence is exemplified by the muscle-specific gene *nautilus* (*nau*), which displayed an unexpected pattern where H3K27ac, H3K27me3, and RNA expression overlap within the muscle lineage (Fig. 3C, D, genome browser shots in Fig. S7A). This finding suggests that Polycomb-mediated silencing is not exclusively used to repress genes directing alternative cell fates; it may also contribute to the restriction of gene expression within a cell lineage. Consistent with this idea, *nau* is only expressed in a subset of muscle precursors to implement their specific differentiation programs (Keller et al., 1998) (Fig. S7B). With our data, we were unable to resolve individual muscle subtypes within the lineage, but it is likely that H3K27me3 is present in one muscle subtype and that H3K27ac and RNA are found in a different type of muscle cell. Other examples of where expression overlaps a high epigenetic potential in a lineage include the genes *HHEX, org-1*, *byn* and *acj6* (Fig. S7C).

**Figure 3.**
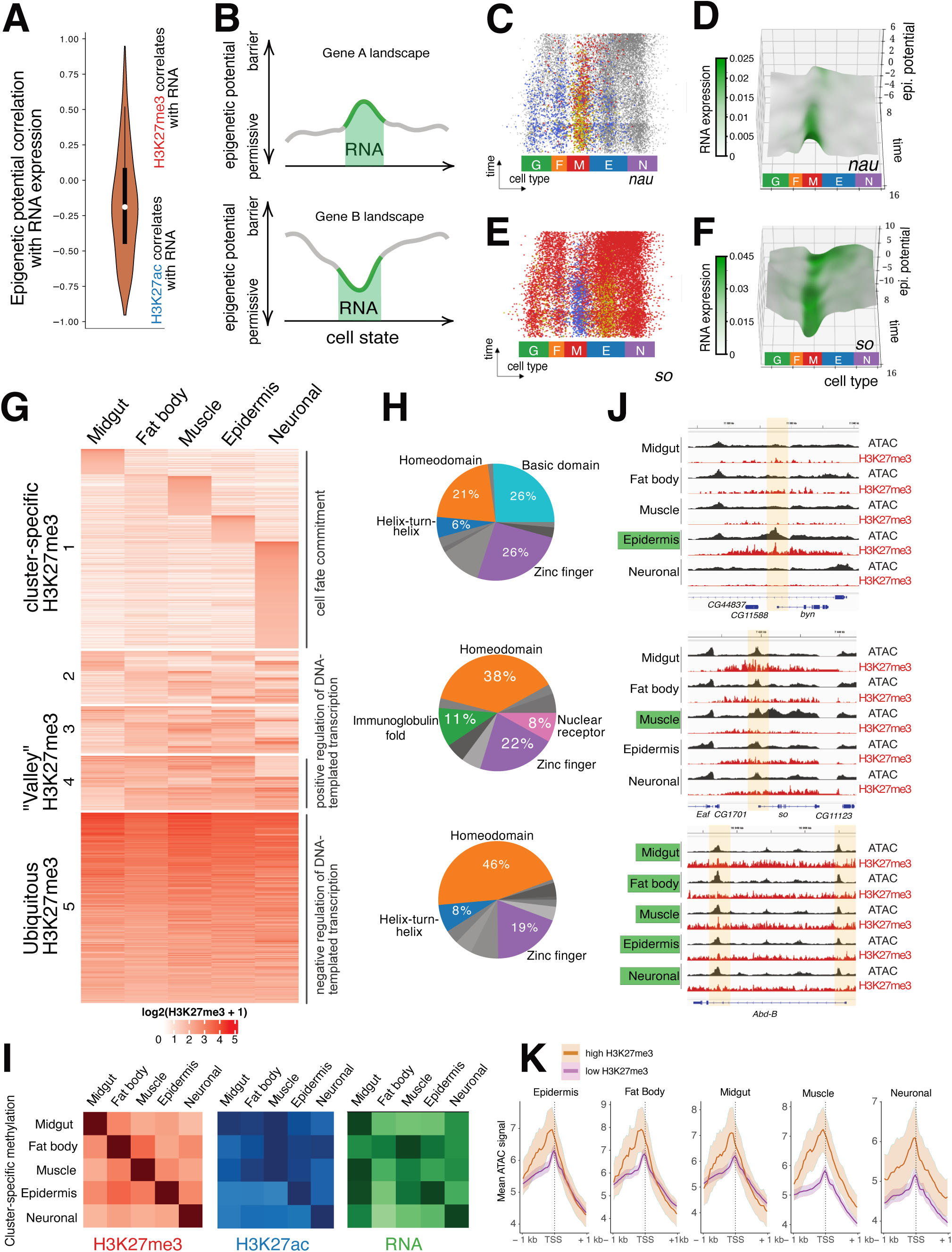
Presence of H3K27me3 in various distributional modes suggest inter- and intra-lineage repression. A. Violin-plot of correlation score (Fisher z-transformed weighted covariances of H3K27ac and H3K27me3) between epigenetic potential and RNA expression on landscapes across expressed genes enriched with both marks (n=3431). B. Depiction of two types of correlation on a time slice of schematic epigenetic landscapes; expression-in-a-valley and expression-in-hills pattern. The area of expression is shown in green, higher epigenetic potential represents putative chromatin barrier, and lower epigenetic potential corresponds to permissive state. C. Cell-time embedding colored by the classified enrichment of H3K27ac/H3K27me3 at bins linked to the muscle-specific gene *nautilus* (*nau)*, indicating acetylation-enriched (blue), methylation-enriched (red), both marks (yellow), or void state (grey color). D. 3D landscape for *nau* colored by inferred RNA expression (x axis: cell type; y axis: inferred developmental time, z-axis: epigenetic potential). Midgut (G), Fat body (F), Muscle (M), Epidermis (E), Neuronal (N). E. Classified enrichment of H3K27ac/H3K27me3 at bins linked to *sine oculis (so)*. F. 3D landscape for *so* colored by inferred RNA expression. G. Heatmap of H3K27me3 signal at bins linked to genes across cell clusters, grouped by number of cell clusters where a gene is methylated, and ordered by the strength of the H3K27me3 signal. The top GO Biological Process term by p-value is shown to the right of the heatmap for the 1^st^, 4^th^, and 5^th^ gene groups. H. Pie charts of transcription factor superclasses found in H3K27me3-marked gene groups (top: first group – cell-type specific methylation; middle: fourth group – valley-shape methylation, bottom: fifth group – ubiquitous methylation).Only the largest classes are shown. Basic domain superclass includes bHLH and bZIP classes. I. Heatmaps of mean H3K27me3 signal, H3K27ac signal and inferred RNA expression at cluster-specific methylated genes (group 1), split by cell type where H3K27 methylation occurs. J. Genome browser snapshots of pseudobulk H3K27me3 and scATAC-seq signals transferred from (Calderon et al., 2022) at representative genes from groupings 1, 4, and 5. Promoters are shown in light yellow. The cell type with major gene expression is highlighted with a green box. K. Metaplot profiles of ATAC-seq pseudobulk signal over TSS of silent genes within a cell cluster split by H3K27-methylated and hypomethylated states.

By contrast, the transcription factor gene *sine oculis* (*so)* is expressed in a low-potential epigenetic valley along the muscle trajectory, characterized by high H3K27me3 in other lineages and high H3K27ac restricted to muscle (Fig. 3F). Although best known as a regulator of eye formation in later stages of development, *so* is reported to function in specifying muscle progenitor cell identity in the embryo (Dubois et al., 2016), and it is expressed exclusively in muscles at this stage based on the (Calderon et al., 2022) scRNA-seq atlas (Fig. 3G). These landscapes demonstrate that RNA expression can both follow an epigenetic valley and co-occur with high levels of H3K27me3.

### H3K27me3 is present in different distributional modes

Another observation related to the co-occurrence of H3K27ac and H3K27me3 was that H3K27 methylation can be cell-type specific, for example in *nau, HHEX*, *byn* and *acj6* (Fig. 3C and Fig. S7A, C). We therefore asked how often H3K27me3 occurs as a cell-type-specific mark overall. We separated genes according to if H3K27me3 was cluster-specific, present in two, three or four cell clusters, or if H3K27me3 was ubiquitous (Fig. 3G), and found that 541 genes were H3K27 methylated in one cell cluster, 153 in two cluster, 122 in three, 161 genes had H3K27me3 in four clusters, and that 308 genes were ubiquitously H3K27 methylated. This shows that the “valley” pattern with H3K27me3 in four out of five cell clusters is observed for a minority (161 out of 1285) of H3K27 methylated genes.

Gene ontology analyses showed that transcription factors were enriched among all types of H3K27me3-decorated genes (Table S4). Basic helix-loop-helix and basic leucine zipper transcription factors were common among cluster specific H3K27me3-decorated genes, but not among ubiquitously H3K27-methylated genes that are instead enriched for homeodomain transcription factor genes (Fig. 3H). The terms cell fate commitment or specification were enriched in cluster-specific and “valley in the hills” H3K27-methylated genes, but not in ubiquitously H3K27me3 genes (Fig. 3G, Table S4). This indicates that cell specification is accompanied by cell-type specific deposition of H3K27me3 over key cell fate determinants. We observed that proximal (+/- 1kb around TSS) Polycomb Response Elements (PREs) are found at 42% of cluster-specific methylated genes, and at 54% of ubiquitously H3K27-methylated genes (Fig. S7D), indicating that non-PRE binding cell-type specific factors are more often involved in recruiting Polycomb proteins to genes with cluster-specific H3K27-methylation. Consistent with this notion, we found that E-box motifs, bound by basic helix–loop–helix transcription factors that are typically associated with cell type–specific promoter activity (Murre, 2019), were enriched among cluster-specific H3K27-methylated genes (Fig S7E, Table S5). By contrast, ubiquitously H3K27me3-decorated genes were most strongly enriched for a GA-rich motif recognized by the ubiquitous factor Clamp (Fig. S7E).

We observed that genes with cluster-specific H3K27me3 in many cases had the highest level of H3K27ac and RNA expression in the same cell cluster, suggesting that the pattern we observed for *nau* (Fig. 3C, D) is common in other cell types (Fig. 3I). Highest expression or acetylation in the same cell cluster as the uniquely H3K27-methylated genes occurred for 20-60% of the genes in different clusters (Fig. S7F). We hypothesize that this cluster-specific mode is required for intra-lineage refinement, restricting expression to a specific cell subset within a permissive lineage.

Why are some genes methylated in multiple cell clusters and others specifically in only one or two? We speculated that this could reflect how accessible the genes are for gene activation in different cell types. By comparing with scATAC-seq data (Calderon et al., 2022), we found several examples where promoters of ubiquitously methylated genes were accessible in all cell types, whereas genes with restricted H3K27me3 were closed or less accessible in cell-types lacking H3K27 methylation (Fig 3J). We compared silent genes with or without H3K27me3 in each of our cell clusters, and found that chromatin accessibility was higher at silent genes with H3K27me3 than silent genes without this mark (Fig. 3K). This indicates that repression of genes lacking H3K27me3 may occur primarily due to nucleosome-mediated occlusion of transcription factor binding, whereas accessible genes may require active deposition of H3K27me3 for silencing.

### Tissue-specific E(z) depletion de-represses tissue-matched targets in the muscle lineage

To test the hypothesis of intra-lineage Polycomb-mediated H3K27me3 repression we performed a cell type-specific RNAi knockdown of the H3K27 methyltransferase Enhancer of zeste, E(z), in mesodermal derivatives using a Mef2-Gal4 driver (Fig. 4A). This Mef2 driver is active in both muscles and fat body (Fig. 4B, S8A). E(z) knockdown with this driver, Mef2>E(z)KD, had a very small effect on embryo hatching and pupariation, but was pharate lethal with close to 100% penetrance (Fig. 4C). We performed single-cell nano-CUT&Tag on 14-16h AEL Mef2>E(z)KD embryos in duplicate, detecting the H3K27me3 and H3K27ac marks (Fig. S8B, C). Although major cell types could be distinguished based on epigenomic profiles, fat body could be separated only after harmonization with the wild-type dataset (Fig.4D, S8D). According to nano-CUT&Tag, H3K27me3 was moderately depleted compared to wild-type in Mef2>E(z)KD embryonic muscle cells, but no global elevation of H3K27ac in muscle was detected (Fig. 4E, S8E). Partial, muscle-specific depletion of H3K27me3 in stage 16 embryos was confirmed by immunostaining (Fig. 4F, S8F). The residual H3K27me3 signal likely comes from maternally loaded E(z) protein (Jones and Gelbart, 1990), which should be unaffected by zygotic knockdown of E(z) mRNA. Genome browser shots illustrate the mesoderm-specific reduction in H3K27me3 over the *E(spl)* and *en-inv* loci (Fig. S8G).

**Figure 4.**
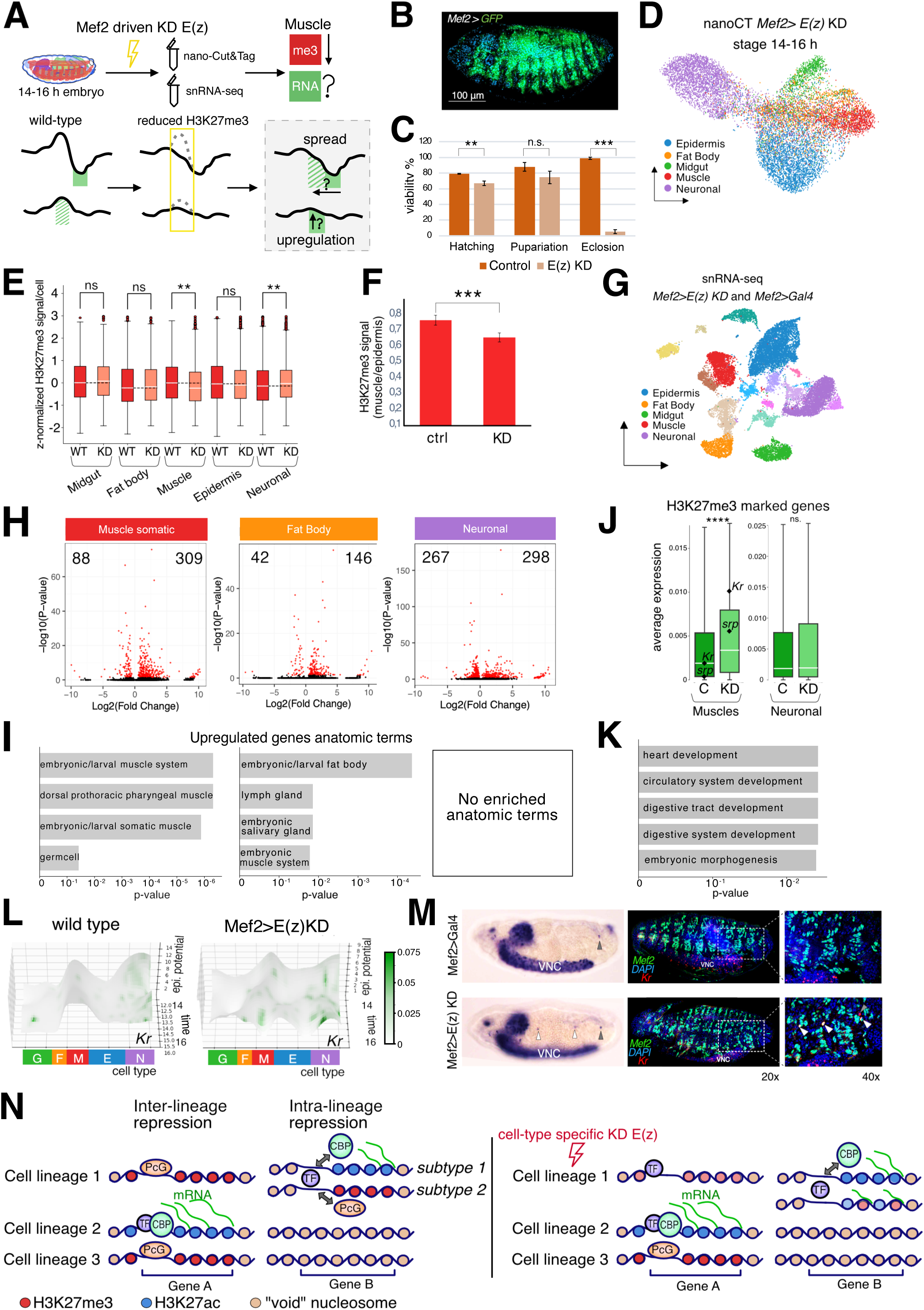
Partial loss of H3K27me3 de-represses tissue-matched targets in Mef2>E(z) knockdown embryos. **A.** Experimental outline of muscle-specific knockdown of E(z), the catalytic subunit of PRC2, and a diagram of two expected outcomes: expansion of RNA expression across lineages and intra-lineage de-repression. **B.** Confocal section of stage 16 embryo expressing Mef2>GFP marking somatic muscles. **C.** Viability assay of Mef2>UAS-E(z)-shRNA (KD) versus Mef2>Gal4 control (C) across embryo hatching, pupa formation, and eclosion. Bars show mean ± s.e.m. from *n* = 3 crosses; ** indicates p-value ≤0.01 and *** indicates p-value ≤0.001, ns – not significant (two-tailed Student’s t-test). **D.** UMAP visualization of cell-bin matrix for bimodal nanoCUT&Tag dataset produced for Mef2>E(z) KD nuclei (14–16 h AEL; 2 biological replicates) after integration with wild-type nanoCUT&Tag dataset; UMAP is colored by annotated cell type. **E.** Boxplots of z-normalized H3K27me3 signal per cell across all bins, comparing Mef2>E(z) KD versus wild-type within each cell type. ** indicates p-value ≤ 0.01 and ns stands for not significant (two-tailed Student’s t-test with Benjamini-Hochberg correction). **F.** Quantification of H3K27me3 fluorescence per nucleus in muscle cells normalized by H3K27me3 intensity in epidermal nuclei from stage 16 embryos (median ± s.e.m.; two-tailed Student’s t-test; *** indicates p-value ≤ 0.001). Five confocal sections of Mef2>E(z) KD stage 16 embryos and four sections of Mef2>GFP stage 16 control embryos were analysed. **G.** UMAP visualization of snRNA-seq co-embedding of integrated Mef2>KD E(z) and Mef2>Gal4 control nuclei from 14–16 h AEL embryos (23,103 nuclei); the embedding is colored by annotated clusters. **H.** Differential expression in representative clusters. Volcano plots for somatic muscle, fat body, and a neuronal cluster. Numbers indicate up/down-regulated genes at p-adj < 0.05. **I.** Enriched anatomic terms among up-regulated genes per cluster (terms ranked by adjusted p-value). **J.** Expression of H3K27-methylated genes (as defined in wild type) in Mef2>GFP control vs Mef2>E(z) KD. Boxplots show mean RNA expression per gene set within Muscle (336 genes) and Neuronal (462 genes) clusters; **** indicates p-value ≤ 0.0001 (two-sided Wilcoxon rank-sum test). **K.** GO Biological Process terms enriched among up-regulated H3K27me3-marked genes in KD muscle (top GO categories). N = 69 genes with p-value < 0.05 and log_2_FC > 0.5. **L.** Epigenetic landscapes for *Kr* gene. The landscape is colored by RNA expression. Left, wild-type 14-16 h AEL embryo. Right, Mef2>E(z) KD 14-16 h AEL embryo. **M.** Validation of *Kr* de-repression. Left, in situ hybridization for *Kr* in Mef2>control and Mef2>E(z) KD; ventral nerve cord (VNC), where *Kr* is expressed at this stage, is indicated. Right, confocal section co-stained for Mef2 (muscle) and Kr protein. White arrowheads: ectopic Kr in muscle; grey arrowhead: endogenous Kr. **N.** Model depicting Polycomb Group (PcG) repression through H3K27me3-marked nucleosomes (red), active genes associated with transcription factors (TF) and H3K27ac nucleosomes (blue) deposited by CBP, and inactive genes with a “void” chromatin state (brown) lacking both H3K27ac and H3K27me3. To the right, the effect of partial lineage-specific H3K27me3 depletion, inducing intra-lineage gene de-repression.

To assess how partial depletion of H3K27me3 affected the transcriptome, we performed single-nucleus RNA-seq (snRNA-seq) in Mef2>E(z)KD and in control Mef2 driver 14-16 h AEL embryos. We profiled 23,103 nuclei in total from two biological replicates for each condition, and the cells were clustered and annotated based on GO and developmental anatomic term enrichment (Fig 4G). Cell types were verified by integration and label transfer from (Calderon et al., 2022) (Fig.4G, Fig. S9A, B, Table S6). Compared to control, the proportion of somatic and visceral muscle, fat body, and plasmatocyte cell clusters were slightly decreased in Mef2>E(z)KD embryos, whereas yolk, salivary gland and one undefined cluster grew in size (Fig. S9C). In mesoderm-derived cell clusters (somatic and visceral muscle, fat body), more genes became significantly up-regulated than down-regulated (Fig. 4H, S9D). Up-regulated genes in the somatic muscle cell cluster were enriched with muscle-related anatomic terms as well as with genes expressed in germ cells, genes up-regulated in visceral muscle were enriched for fat body genes, while fat body up-regulated genes were annotated as being expressed in fat body, lymph gland, salivary gland and muscle (Fig. 4I, Table S7). By contrast, no anatomic term was enriched in up-regulated genes in neurons.

Because these deregulated genes can be both direct and indirect targets of the knockdown, we focused on genes marked with H3K27me3 in muscle. These genes were significantly upregulated in Mef2>E(z)KD muscle as a group (Fig. 4J, Table S8). The H3K27me3 marked up-regulated genes were enriched with GO-terms related to heart and circulatory system development, as well as digestive tract development and morphogenesis (Fig.4K).

H3K27me3-decorated genes in the neuronal clusters were not affected as a group (Fig. 4J). Many up-regulated genes in muscle were slightly less H3K27-methylated than unaffected genes, and gained H3K27ac (Fig. S10A, B). Thus, snRNA-seq revealed that the partial loss of H3K27me3 primarily led to de-repression of genes that are normally expressed in the same lineage. These upregulated genes were significantly enriched for tissue-matched functional terms.

To further validate this result, we examined expression of the *Krüppel* (*Kr*) gene. The *Kr* transcription factor is strongly expressed in brain and ventral nerve cord at stage 16 (Isshiki et al., 2001; Romani et al., 1996), but also determines specific muscle identities at stage 11-15 (Ruiz-Gomez et al., 1997). *Kr* was strongly methylated in muscle (Fig. 4L, Fig. S10C). We therefore reconstructed its chromatin landscape, comparing wild-type and 14-16h Mef2>E(z)KD datasets, illustrating the reduced level of H3K27 methylation along the muscle trajectory in Mef2>E(z)KD embryos (Fig. 4L, right panel, genome browser shot shown in Fig. S10D). Using whole-mount in situ hybridization and co-immunostaining for Kr and Mef2, we detected ectopic Kr expression in a few muscle cells in Mef2>E(z)KD embryos (Fig. 4M). Together, these results support the conclusion that even a partial decrease in H3K27me3 can lead to de-repression of tissue-specific genes.

Another H3K27-methylated gene that was up-regulated in muscle is *serpent* (*srp*) (Fig. 4J). This GATA transcription factor is most strongly expressed in fat body in late stage embryos, but has also earlier functions in hematopoiesis and gut differentiation (Hayes et al., 2001; Rehorn et al., 1996). By contrast, partial reduction of the epigenetic barrier did not cause ectopic expression of epidermal cell fate determinants such as *vvl* or *grh* in the muscle lineage (Table S10). Taken together, these results suggest that partial loss of H3K27me3 primarily causes de-repression of genes expressed in the same lineage, whereas restriction of cell fate determinants to separate cell lineages remain intact.

## Discussion

Our study provides evidence of how the co-dynamics of H3K27ac and H3K27me3 shape the developing chromatin landscape during Drosophila embryogenesis, offering a visual foundation for the epigenetic barriers that may contribute to Waddington’s conceptual model. We introduce a novel analytical framework - a chromatin state-based epigenetic landscape - and reveal an intra-lineage role for H3K27me3 in restricting gene expression within committed cell types.

Cell state transitions are governed by gene regulatory networks, and quantitative reconstruction of the Waddington landscape has been attempted recently based on modelling or on single-cell transcriptomic data (Cislo et al., 2025; Han et al., 2025). To what extent chromatin state contributes to the energy barriers that restrict cell state transitions is not known, but chromatin-based memory has been proposed to allow cell states to remain stable (Bell et al., 2024). A more comprehensive understanding of chromatin state in cell fate choice and maintenance is warranted, but histone modification profiles in individual cells and cell types lag behind the numerous scRNA-seq datasets that are available (Liu et al., 2025).

Although single-cell profiling of histone modifications has been possible for some time (Ai et al., 2019; Bartosovic et al., 2021; Grosselin et al., 2019; Ku et al., 2019; Wang et al., 2019; Wu et al., 2021), simultaneous mapping of multiple histone marks in single cells has only recently become feasible (Bartosovic and Castelo-Branco, 2023; Gopalan et al., 2021; Meers et al., 2023; Stuart et al., 2023; Xiong et al., 2024). Here, we employ nano-CUT&Tag to generate an organismal and cell-specific histone modification atlas for H3K27ac and H3K27me3 at the same time.

By simultaneously mapping H3K27ac and H3K27me3, we established a metric for epigenetic potential. Our cell type-time embedding allowed us to visualize how this potential changes along developmental trajectories, directly modelling the ‘valleys’ and ‘hills’ of a chromatin landscape for individual genes. Although chromatin landscapes have been generated for mouse development (Gorkin et al., 2020; Yu et al., 2025), our analysis of epigenetic potential provides a direct visualization of the chromatin barriers that contribute to cell-specific gene expression. We observed a global trend toward increased restriction—more cell type-specific H3K27ac and a gain of H3K27me3 over time. In the Drosophila embryo, increased restriction over developmental time is also observed for transcription factor expression (Adryan and Teichmann, 2010), suggesting that transcription factors and chromatin state cooperate in generating cell specific gene activity. The visualization of gene expression following the low-potential valleys reinforces the idea that chromatin state acts as a constraint on the transcriptome.

A function for PRC2 and H3K27me3 in cell lineage restriction has also been described in mammalian systems, but has mostly focused on its role in embryonic and adult stem cells (Brand et al., 2019). For example, Polycomb silencing restricts differentiation of naïve pluripotent stem cells towards the trophoblast lineage (Kumar et al., 2022; Zijlmans et al., 2022), prevents differentiation of primed embryonic stem cells (Collinson et al., 2016; Shan et al., 2017), increases lineage fidelity during anterior endoderm differentiation (Holzenspies et al., 2025), and represses undesirable differentiation programs in epidermal and intestinal progenitors (Chiacchiera et al., 2016; Ezhkova et al., 2009). In brain organoids, inhibiting EZH2 and H3K27me3-deposition results in aberrant cell identity acquisition and a shift of neural progenitor cell proliferation toward differentiation (Ditzer et al., 2025; Zenk et al., 2024).

However, our findings contradict the view of H3K27me3 as a primarily broad, inter-lineage repressive mark (Fig. 4N). We demonstrate that H3K27me3 exists in distinct distributional modes: broad/ubiquitous repression (the canonical role of PRC2 in silencing alternative cell fates) and cluster-specific repression. We propose that the cluster-specific mode is required for intra-lineage refinement. Genes in this category often display a nucleosome-occupied “void” state in non-permissive tissues (Schwartz et al., 2010), but require active PRC2 repression within the permissive lineage to prevent full transcriptional activation in every cell, thus restricting expression to a specific cell subset and fine-tuning heterogeneity.

Cell-type-specific E(z) knockdown provided functional evidence for this hypothesis. The partial loss of H3K27me3 de-repressed genes that are normally expressed in the same tissue, e.g., *Kr* in muscle cells. The tissue-matched nature of the upregulated genes supports that H3K27me3 acts to counteract existing activation cues within the differentiating muscle lineage, thereby stabilizing and restricting the final transcriptional program.

In conclusion, our study reframes H3K27me3 not only as a barrier between cell fates but also as a critical tuner of expression heterogeneity that contributes to the final topography of the epigenetic landscape within committed lineages.

## Materials and Methods

### Drosophila stock maintenance

*yw; PCNA-eGFP*, a kind gift of Eric Wieschaus (Blythe and Wieschaus, 2016), served as a control (wild-type) line for the nano-CUT&Tag experiments. To perform knockdown experiments, we crossed *yw; Mef2-GAL4* (Bloomington 27390) males with *y; UAS-E(z) RNAi* (*y^1^ sc v^1^ sev^21^; P{y^+t7.7^ v^+t1.8^=TRiP.HMS00066}attP2*, Bloomington 33659) virgin females. *yw; Mef2-GAL4* served as a control line for the experiments with E(z) knockdown. For Mef2-driven GFP expression we crossed *w; UAS-GFP.nls* (Bloomington 4775) males with *yw; Mef2-GAL4* virgin females. For combined expression of GFP and E(z) RNAi under Mef2 control, we established the stock *w; UAS-GFP; UAS-E(z)* RNAi flies, and crossed the males from this stock with *yw; Mef2-GAL4* virgin females.

Stocks were kept on potato mash-agar food and maintained at 25°C with a 12-h light/dark cycle. Embryos were collected on apple juice plates supplemented with fresh yeast and aged at 25°C for specific time ranges dependent on the specific experiment which is detailed in the relevant methods section. Plates containing embryos collected for the first 2 h each day were discarded to avoid contamination by older embryos withheld by females.

### Viability assay

We collected embryos with Mef2 driven E(z) knockdown and control Mef2-GAL4 ones for four hours and kept them for 36 h at 25°C. After that we assessed the number of empty chorions and unhatched embryos, to determine hatching rates. From each plate we placed 50 first instar larvae into vials. After one week we counted the number of pupae to estimate the pupariation rate (number of pupae divided by 50). Two weeks after placing first instar larvae into vials we determined eclosion rates by dividing the number of empty pupae by total number of pupae. At least two replicates were performed for each experimental genotype.

### nanoCUT&Tag

#### Drosophila embryo nuclear extraction

For nanoCUT&Tag (nanoCT), embryos were collected on apple juice agar plates supplemented with yeast for 2 hours and aged an additional 8 hours or 14 hours, following two sequential two-hour pre-collections. Collected embryos were dechorionated in 5x diluted bleach, rinsed thoroughly in embryo wash buffer (PBS, 0.1% Triton X-100) and transferred to HB buffer (15 mM Tris-HCl pH 7.4, 0.34 M sucrose, 15 mM NaCl, 60 mm KCl, 0.2 mM EDTA, 0.2 mM EGTA, freshly added protease inhibitor cocktail (cOmplete ULTRA tablets, Roche). For nuclear extraction, we followed the protocol of (Calderon et al., 2022). In short, we used a Dounce homogenizer 20 times with a loose pestle and 10 times with a tight pestle. Then, we filtered nuclei through two Miracloth (Millipore, #475855) layers to another Eppendorf LoBind tube and spun for 5 min 2000 rcf. The pellet was washed in 1 ml of HB buffer and spun again. The washed pellet was resuspended in PBTB buffer (1x PBS, 0.1% Triton X-100, 5% BSA). Nuclei were dissociated by passing them through 19G and 22G needles in a 2ml syringe and filtered through 20 µm nylon net (Millipore #NY2004700).

After that, nuclei were stained with Trypan Blue 0.4% (HyClone, #SV30084.01) and inspected for the presence of clumps or debris under a bright-field microscope and manually counted.

#### Single-cell nanoCT

We followed the nano-CT protocol with minor modifications (Barcenas-Walls et al., 2024; Bartosovic and Castelo-Branco, 2023). NanoCT experiments were performed in two independent biological replicates for each time point or condition. For each sample 400K nuclei were centrifuged for 10 min at 700g at 4°C and the supernatant removed. The nuclear pellet was resuspended in 200 μL antibody buffer (20 mM HEPES pH 7.5, 150 mM NaCl, 2 mM EDTA, 0.5 mM spermidine, 0.05% digitonin, 0.01% NP-40, 1x protease inhibitors and 2% bovine serum albumin (BSA)). Nuclei were centrifuged for 5 min at 700g and resuspended in 100 μL of antibody buffer. Each sample was aliquoted in 0.5 ml Eppendorf LoBind tubes, and 1 μL of primary antibody against H3K27me3 (Ab6002), 1 μL of primary antibody against H3K27a. (Abcam, ab177178), 1 μL of anti-mouse nano-Tn5, 1 μL of anti-rabbit nano-Tn5 (Protein Science Facility, Karolinska Institutet, Stockholm) with different barcoded ME-P5 oligonucleotides were added into each tube.

The samples were gently resuspended and incubated at 4°C on a roller overnight. After overnight incubation, the nuclei were centrifuged at 700g for 10 min, the supernatant was discarded and washed twice with Dig-300 wash buffer (20 mM HEPES pH 7.5, 300 mM NaCl, 0.5 mM spermidine, 0.05% digitonin, 0.01% NP-40, 2% BSA, and 1× protease inhibitors). The nuclei were resuspended in 200 μL of tagmentation buffer (20 mM HEPES pH 7.5, 300 mM NaCl, 0.5 mM spermidine, 0.05% digitonin, 0.01% NP-40, 2% BSA, 1× protease inhibitors, and 10 mM MgCl_2_) and incubated for 1 h at 37°C with resuspending the nuclei by pipetting after 30 min. The reaction was stopped by addition of 200 μL of Stop buffer (200 μl 1× DNB from 10x scATAC-seq kit, supplemented with 2% BSA and 25 mM EDTA) and pipetted three times. The nuclei were sedimented by centrifugation for 10 min at 700g, the supernatant discarded and nuclear pellet was resuspended in 200 μL of 1X DNB+2% BSA buffer. A second centrifugation at 10 min 700g was performed, and the supernatant has been partially removed from the nuclear pellet to leave approximately 10 μL. After the addition of 10 μL of 1X DNB+2% BSA buffer, nuclei were resuspended and manually counted using a solution consisting of 2 μL of nuclei suspension and 8 μL of Trypan Blue 0.4%.

#### Single-cell indexing and library prep

Single-cell indexing was performed using Chromium Next GEM Single Cell ATAC Library & Gel Bead Kit v2.

We mixed 16000 nuclei (up to 8 μL of final nuclei suspension from the last step) with 7 μL 10x ATAC Buffer B, 56.5 μL Barcoding Reagent B, 1.5 μL Reducing Agent B and 1 μL Barcoding Enzyme. Seventy microliters of the master mix and recovered nuclei were loaded on Chromium Next GEM Chip H, followed by single-cell partitioning using the Chromium controller and GEM incubation according to the manufacturer’s instructions (Chromium Next GEM Single Cell ATAC Reagent Kits v2 (Steps 2.0–2.5)). After completing the GEM incubation in the thermal cycler, the sample was recovered by post-GEM incubation cleanup with Dynabeads MyOne SILANE and SPRIselect beads according to Chromium Next GEM Single Cell ATAC Reagent Kits v2. (Steps 3.0–3.2).

After elution from SPRIselect beads, 40 μL samples were recovered and used for a second round of tagmentation with Tn5-ME-B (P7) transposome. The following reaction mix was used: 38 μl template DNA, 50 μl 2× TD buffer (20 mM Tris pH 7.5, 10 mM MgCl2, and 20% dimethylformamide), 0.2 μl Tn5 (loaded with ME-B only, and previously titrated) per 10 ng of template DNA and nuclease-free water up to 100 μl, and incubated for 30 min at 37 °C in a thermocycler. After that, samples were purified with DNA clean and concentrator-5 kit (Zymo, #D4014) and eluted in 40 μL of elution buffer. The purified DNA was used as template for the final PCR amplification, following the Chromium Next GEM Single Cell ATAC Reagent Kits v2 (Step 4.1) (40 μL sample, 50 μL AMP Mix, 7.5 μL SI-PCR primer B and 2.5 μL individual Single index N Set A primer) and amplified using standard 10x scATAC-seq PCR amplification program (1. 72°C 5 min, 2. 98 °C 45 s; 3. 98 °C 20 s; 4. 67 °C 30 s; 5. 72 °C 20 s; 6. 72 °C 1 min; 7. 4 °C hold) for 16 cycles. The amplified product was purified with SPRIselect reagent (Beckman-Coulter, #B23317) using 0.4X and 1.2X two-sided purification according to the Chromium Next GEM Single Cell ATAC Reagent Kits v2 (Step 4.2). The library concentration and size distribution were assessed with Qubit dsDNA HS Assay Kit (ThermoFisher) and Bioanalyzer High-Sensitivity DNA kit (Agilent). The libraries were sequenced on Illumina Novaseq X (1.5B) NovaSeq SP-100 using custom read1 (R1_seq GCGATCGAGGACGGCAGATGTGTATAAGAGACAG) and custom index2 (I2_seq CTGTCTCTTATACACATCTGCCGTCCTCGATCGC) primers with the following read lengths 36-8-48-36 (R1-I1-I2-R2).

### Single-nucleus RNA sequencing

For snRNA-seq, embryos were collected on apple juice agar plates supplemented with yeast for 2 h and aged for an additional 14 h, following two sequential 2 h pre-collections. Embryos were dechorionated in diluted bleach, thoroughly rinsed in embryo wash buffer (PBS, 0.1% Triton X-100), and transferred to HB buffer (15 mM Tris-HCl, pH 7.4, 0.34 M sucrose, 15 mM NaCl, 60 mM KCl, 0.2 mM EDTA, 0.2 mM EGTA, supplemented fresh with protease inhibitor cocktail; cOmplete ULTRA tablets, Roche).

Nuclei were isolated using the same procedure as in the nanoCT experiment described by Calderon et al. (2022). Briefly, embryos were homogenized, the homogenate was filtered through Miracloth, and nuclei were pelleted by centrifugation for 5 min at 2,000 × g. The pellet was washed once in 1 ml HB buffer and centrifuged again under the same conditions. The washed nuclei were then resuspended in PBTB buffer (1× PBS, 0.1% Triton X-100, 0.04% BSA). To dissociate nuclei, the suspension was passed sequentially through 19G and 22G needles using a 2 ml syringe, and then filtered through a 20 μm nylon mesh. Nuclei were inspected by bright-field microscopy and manually counted.

A total of 20,000 nuclei were loaded for snRNA-seq library preparation using the MGI DNBelab C Series High-throughput Single-cell RNA Library Kit, according to the manufacturer’s instructions. Libraries were sequenced on a DNBSEQ-G400RS instrument using the High-throughput Sequencing Reagent Set. The experiment was performed with two biological replicates per condition.

For snRNA-seq, embryos were collected on apple juice agar plates supplemented with yeast for 2 h and aged for an additional 14 h, following two sequential 2 h pre-collections. Embryos were dechorionated in diluted bleach, rinsed in PBS + 0,1% Triton X-100 and transferred to HB buffer (15 mM Tris-HCl pH 7.4, 0.34 M sucrose, 15 mM NaCl, 60 mm KCl, 0.2 mM EDTA, 0.2 mM EGTA, freshly added protease inhibitor cocktail; cOmplete ULTRA tablets, Roche). Nuclei were isolated using the same procedure as in the nanoCT experiment.

Briefly, embryos were homogenized, the homogenate was filtered through Miracloth, and nuclei were pelleted by centrifugation for 5 min at 2,000 × g. The pellet was washed once in 1 ml HB buffer and centrifuged again under the same conditions.

The washed nuclei were then resuspended in PBTB buffer (1× PBS, 0.1% Triton X-100, 0.04% BSA). To dissociate nuclei, the suspension was passed sequentially through 19G and 22G needles using a 2 ml syringe, and then filtered through a 20 μm nylon mesh. Nuclei were inspected by bright-field microscopy and manually counted.A total of 20,000 nuclei were loaded for snRNA-seq library preparation using the MGI DNBelab C Series High-throughput Single-cell RNA Library Preparation Set V3.0 (TaiM 4), according to the manufacturer’s instructions. Libraries were sequenced on a DNBSEQ-G400RS instrument using the High-throughput Sequencing Reagent Set. The experiment was performed with two biological replicates per condition.

### Bioinformatic analyses

#### Nano-CUT&Tag data processing and integration

##### NanoCT preprocessing

The nano-CUT&Tag (nanoCT) sequencing FASTQ files were processed with nanoscope, github:bartosovic-lab/nanoscope using GTACGGTT as a modality barcode for H3K27ac and GGCTCTGA for H3K27me3, which created a fragment file. We then created a count matrix by assigning all fragments into 5000 bp genomic bins. Cells were filtered to have >20 fragments for either modality and to be below the 99^th^ percentile based on the fragment number to remove potential doublets. Features were filtered to a minimum of 1 fragment. To embed our data, we used the multi-view spectral algorithm from SnapATAC2 (n_components = 25), which combines information from both modalities (Zhang et al., 2023). We removed one (almost always first) principal component (PC), where the PC correlates |ρ| > 0.6 with H3K27me3/H3K27ac summed counts. We reasoned that this PC likely contains information about read depth per cell. UMAP was created on the spectral embedding by including all components with mostly default parameters and n_neighbors = 15. For plotting the H3K27ac and H3K27me3 signal on the UMAP, we normalized counts by total counts per cell barcode and transformed the data by log10(x+1). We performed Leiden clustering with a resolution 0.5 capturing the major cell types in the embryo.

##### Cluster annotation

Cell clusters were annotated independently for the 8–10 h AEL, 14–16 h AEL, combined time points after integration with scATAC, and 14–16 h AEL E(z) knockdown datasets using Gene Ontology Biological Process (GO BP) terms. Cluster-specific H3K27ac marker gene statistics were calculated with a likelihood ratio test adapted from SnapATAC2, in which each cluster was compared against all remaining cells. Marker genes were defined by log2FC > 0.2 and p-value < 0.05. GOATOOLS-based GO term enrichment analysis, as well as a custom anatomical-term enrichment analysis, was then performed on top 200 marker genes filtered by p-value.

To refine and validate cluster annotation, we generated a custom GO-like namespace of anatomical terms derived from gene expression localization annotations in the BDGP in situ hybridization database. For annotation of the 8–10 h AEL dataset, we used BDGP expression terms from “range 5,” corresponding to embryonic stages 11–12. For the 14–16 h AEL dataset, we used “range 6” terms, corresponding to stages 13–16. In both cases, we manually introduced higher-level umbrella terms to enable term aggregation through the propagate_counts option in GOATOOLS. For example, the terms “embryonic central brain glia” and “embryonic central brain neuron” were both assigned to the umbrella term “brain.” The annotation term “strong ubiquitous” was excluded. This yielded a total of 12,763 gene–term annotations across 127 terms for range 5, and 20,508 gene–term annotations across 139 terms for range 6. GOATOOLS (Klopfenstein et al., 2018) was then applied to this custom namespace to compute enrichment statistics based on BDGP in situ expression data.

Following identification of cluster-specific acetylation markers, enriched GO BP terms and anatomical terms were compiled for each cluster and used to guide annotation. Final annotations were further validated by comparison with labels transferred from the scATAC-seq dataset using k-nearest-neighbor classification on the GLUE embedding (n_neighbors=1; scikit-learn).

The BDGP-derived umbrella anatomical terms were also used to define gene modules for UMAP projection. For each term, the corresponding module score was calculated as the summed H3K27ac signal across all genes assigned to that umbrella category.

##### nanoCT pseudobulk browser track generation

Pseudobulk bigwig files for each major cluster per time point were generated using pybedtools (https://daler.github.io/pybedtools/) by normalizing the genomic fragment distribution by total fragment number in a cluster.

##### NanoCT integration with scATAC-seq and scRNA-seq

To compare our nanoCT data with published scATAC-seq and scRNA-seq datasets, we downloaded raw count matrices from the DEAP resource (Calderon et al., 2022). For scATAC-seq, we used the time windows relevant to our analysis, namely exp1_8-12h and exp1_12-16h. For scRNA-seq, we used the 6-10h, 8-12h, 10-14h, 12-16h, and 14-18h collections.

To integrate nanoCT data with scATAC-seq or scRNA-seq data, we used GLUE (Cao and Gao, 2022). We first integrated the scRNA-seq and scATAC-seq datasets using GLUE with latent_dim = 20, a negative binomial (NB) model, and a guidance graph linking each promoter to genomic bins within a ±20 kb window. After integration, this graph was pruned on the basis of feature-embedding correlations to generate a regulatory graph connecting bins to genes (see Extended Methods 1.1 for the promoter-distance weighting function). For each gene, the bin overlapping the promoter region was always included, using the most upstream promoter in cases with alternative promoters.

To convert the scATAC-seq bin-by-cell matrix into a gene-by-cell matrix, we used the GLUE-derived gene–bin regulatory graph. Specifically, the adjacency matrix of this graph was multiplied by the bin-by-cell matrix, and the resulting values were normalized by the number of bin connections assigned to each gene.

We then integrated nanoCT with scATAC-seq in GLUE using latent_dim = 20 and a NB model. For this integration, the guidance graph was constructed by linking genomic bins across the two datasets, such that each scATAC-seq genomic locus was connected to two nanoCT loci, one corresponding to each chromatin modality.

##### Shannon entropy

Shannon entropy was calculated separately for the 8-10 h and 14-16 h nanoCT datasets after depth normalization of per-gene fragment counts. Entropy was computed across five cell types using scipy.stats.entropy with base = 2 (Virtanen et al., 2020). For each modality, we selected the top 200 cluster-specific genes ranked by p-value.

##### Epigenetic signal profile over time

To visualize changes in chromatin signal over developmental time, we generated time series of single-cell epigenetic signal for each modality and gene using neural-network age inferred from the integrated scATAC-seq dataset (Calderon et al., 2022). Signals were binned into 1-hour intervals across the 8–16 h developmental window and smoothed using a one-dimensional Gaussian filter (σ = 3). For visualization, the acetylation signal was scaled by a constant. We do not show time points in between 8 and 16 hours since our collection windows were gapped, and the continuous changes in epigenetic marks result from the integration process, therefore making impossible to describe the precise time of changes.

To quantify increases and decreases in H3K27ac and H3K27me3 between early and late time points, we applied a likelihood ratio test adapted from SnapATAC2, comparing each late time-point cell type with its corresponding early time-point counterpart. Genes were classified as dynamic if they satisfied the following criteria: absolute log_2_FC > 0.2, p-value < 0.1, and mean signal > 0.6 for H3K27me3 or > 0.7 for H3K27ac.

##### Correlation of epigenetic marks between cell types

To assess similarity in chromatin profiles across cell types, we calculated Pearson correlation matrices separately for 8–10 h AEL and 14–16 h AEL cells for H3K27ac and H3K27me3.

Epigenetic signal was depth normalized prior to analysis. Features were filtered by mean signal, using thresholds of > 0.5 for H3K27ac and > 2.0 for H3K27me3. This resulted in 11,726 genes for 8–10 h H3K27ac, 772 genes for 8–10 h H3K27me3, 11,636 genes for 14–16 h H3K27ac, and 709 genes for 14–16 h H3K27me3.

### Generating a chromatin landscape

To generate a cell-time embedding, we defined two coordinates: developmental time and cell type. Developmental time was obtained directly from the integrated scATAC-seq dataset, in which cell age had previously been estimated using a neural-network-based predictor (Calderon et al., 2022). These age values were transferred to nanoCT cells by k-nearest-neighbor regression (n_neighbors = 1).

To generate a one-dimensional continuous cell-type axis, we aimed to preserve lineage integrity and developmental relationships while allowing cell identity to vary smoothly along a single coordinate. We therefore first focused on late embryonic cells, reasoning that terminal states provide a stable reference for lineage organization. Cells with predicted neural-network age > 14 were selected, and a low-resolution Leiden clustering (resolution=0.09) was performed after construction of a k-nearest-neighbor graph. The resulting clusters were then arranged into a hierarchical dendrogram by iterative fusion based on kNN adjacency (Dreveton et al., 2025). This procedure produced an ordered hierarchical scaffold that defined the cell-type axis after dendrogram ordering by kNN adjacency (Extended Methods 2.1). The resulting relationships between cell types were biologically correct.

Next, we generated metacells across the full scATAC-seq dataset spanning 8 to 16 h AEL using high-resolution Leiden clustering (resolution=4). Each metacell was assigned an x-coordinate according to its kNN adjacency to the previously defined cell-type scaffold (Extended Methods 2.1), and a y-coordinate corresponding to the mean neural-network age of the cells within that metacell. Individual cells were then embedded according to their specific age and the inferred cell-type coordinate of their assigned metacell. To improve local structure while preserving the intended global organization, we performed a limited number of UMAP iterations (n = 3). The resulting embedding was termed the cell-time embedding.

To generate the three-dimensional epigenetic landscape, we defined the z-coordinate as epigenetic potential, calculated as the difference between H3K27me3 and H3K27ac signal. Because the two modalities could not be directly normalized in a straightforward manner, the acetylation signal was scaled by a genome-wide constant chosen to provide biologically meaningful contrast when visualizing representative genes of interest. A regular two-dimensional grid was then generated using numpy.mgrid (Harris et al., 2020), spanning the same coordinate range as the cell-time embedding. For each grid cell, the signal value was calculated as the mean of all cells falling within that bin. The resulting surface was smoothed using a modified median filter followed by a Gaussian filter to obtain the final landscape representation (Extended Methods 2.2).

To project RNA expression onto the three-dimensional epigenetic landscape, we used kNN-based regression on the GLUE-integrated embedding to predict the cell-time coordinates of cells from the scRNA-seq dataset.

### Cell classification based on chromatin marks

To assign each gene to a predominant chromatin state within the cell-time embedding, we developed a binary classification scheme that distinguished whether a gene was primarily associated with H3K27ac, H3K27me3, or showed comparable probabilities of carrying either mark. For each gene and modality, we sought to determine whether the mark was present or absent in individual cells.

Because sparse single-cell chromatin data are strongly influenced by local neighborhood structure, we designed the classification to operate on the data manifold rather than on individual observations alone. We therefore first generated a normalized score matrix that incorporated local cell-community information. Cell communities were defined using a random cutting algorithm on the k-nearest-neighbor graph (Extended Methods 3.1.1). These randomly generated communities were then used to compute community-level scores for each gene and chromatin modality.

Binary presence/absence calls for each mark were obtained by thresholding this score matrix with two free parameters (Extended Methods 3.2). To determine suitable parameter values, we examined classification outputs for 10 genes of interest across a regularly sampled parameter grid and selected a=0.07 and b=1.9 for downstream analyses.

### Correlation between RNA expression and chromatin landscape

To quantify the relationship between RNA expression and histone marks, we used the landscape grid data generated after landscape embedding. Because a standard Pearson or Spearman correlation applied directly to the grid data would treat all regions equally, including areas with little or no signal, we instead designed a weighted correlation framework that emphasizes informative regions of the landscape. For each gene, grid values were first normalized by Winsorization followed by min–max scaling. We then calculated weighted covariances between RNA expression and H3K27ac, and between RNA expression and H3K27me3. The weights were defined as the squared summed signal of each nanoCT modality with RNA expression after normalization, thereby strongly upweighting grid regions with high chromatin mark or RNA signal and sharply downweighting regions with little or no signal.

To estimate the relationship between epigenetic potential and RNA expression, we combined the correlations of H3K27ac with RNA and H3K27me3 with RNA. These correlation values were first Fisher z-transformed to permit additive and subtractive operations. The final epigenetic potential–RNA correlation score was then defined as (also see Extended Methods 5): *corr_epi−RNA_* = tanh(*z_H_*_3*K*27*me*3*−*_*z_H_*_3*K*27*ac*_)

, where z_H3K27me3_ and z_H3K27ac_ are the Fisher-transformed correlations of methylation and acetylation with RNA, respectively. Genes were included based on the following thresholds: mean H3K27me3 signal > 0.15, mean H3K27ac signal > 0.15, mean combined epigenetic signal (H3K27me3 + H3K27ac) > 1.5, and mean RNA expression > 0.0005, resulting in a total of 3,431 genes retained for analysis.

### NanoCUT&Tag data analysis from E(z) knockdown embryos

#### Global chromatin signal normalization in the knockdown dataset

To visualize the effect of Mef2>E(z) knockdown on H3K27 methylation levels in mesoderm-derived cell types, we aimed to compare fragment counts across cell types. However, after construction of the cell x bin matrix, the Mef2>E(z)KD dataset showed a different fragments-per-cell distribution, both in mean and variance, relative to wild type. To enable comparison between conditions, we therefore included all wild-type (n=10,450) and Mef2>E(z)KD (n=10,111) nuclei and performed batch normalization. Fragment counts were then summed across all bins for each cell, log2-transformed, and z-normalized within each condition (Extended Methods 6.1).

#### Differential gene-level analysis of chromatin marks in Mef2>E(z)KD

To compare per-gene H3K27me3 and H3K27ac levels between wild type and Mef2>E(z)KD, we applied a likelihood ratio test implemented in SnapATAC2. Because the knockdown dataset displayed a different distribution of ranked fragments per gene relative to wild type, which we considered likely to be technical in origin, we rescaled the knockdown distribution to match the wild-type distribution using a rank-dependent coefficient (Extended Methods 6.2).

#### Epigenetic landscapes for E(z) knockdown experiment

To generate epigenetic landscapes for the knockdown experiment, we first integrated the two nanoCT datasets, wild type and Mef2>E(z)KD (14–16 h AEL, two biological replicates), using Harmony. We then projected Mef2>E(z)KD cells onto the existing wild-type cell-time embedding by k-nearest-neighbor regression (n_neighbors=1).

To project gene expression from the snRNA-seq experiment onto these landscapes, we first integrated wild-type nanoCT with the Mef2>GFP control snRNA-seq dataset using GLUE, following the same procedure as described for the scATAC-seq/scRNA-seq integration. In parallel, Mef2>E(z)KD snRNA-seq was integrated with Mef2>GFP snRNA-seq using Harmony. After these integrations, RNA expression values were projected onto the three-dimensional landscapes using k-nearest-neighbor regression.

### Gene clustering by H3K27me3 distribution mode

We considered only genes with H3K27me3 signal > 2 in at least one cell cluster. Because our aim was to capture the specificity of H3K27me3 distribution across cell types, we emphasized relative enrichment rather than absolute signal. A gene was classified as H3K27-methylated in a given cell cluster if its signal in that cluster was either the highest across all clusters or within twofold of the highest signal. Conversely, if the signal in a cluster was less than half of the maximum signal observed for that gene across all clusters, the gene was classified as hypomethylated in that cluster, even when the absolute signal exceeded 2.

Genes were then grouped according to the number of cell clusters in which they were classified as H3K27-methylated. Within each group, genes were first ordered by the combination and order of cell clusters in which methylation was detected, and then by signal strength.

### Transcriprion factor superclasses, PRE and Motif enrichment analysis

For gene sets displaying ubiquitous, valley-like, or cluster-specific H3K27me3 patterns, transcription factor genes were assigned to superclasses based on the FlyBase gene group dataset (available at https://s3ftp.flybase.org/releases/current/precomputed_files/genes/index.html).

For the same gene sets, we extracted promoter coordinates for all genes, defined as TSS ± 1 kb. Promoter overlap with Polycomb Response Elements (PREs) was identified using coordinates from PRE Mapper and the bedtools intersect tool.

Motif enrichment analysis was performed using SEA from the MEME Suite, with all remaining genes used as the control set (Bailey and Grant, 2021). Motif position weight matrices were obtained from the redundant JASPAR collection (Ovek Baydar et al., 2026). Motifs with E-value ≤ 1 were retained.

### snRNA-seq data analysis

#### Read processing and count matrix generation

Read demultiplexing was performed in real time by DNBSEQ-G400RS software. The resulting FASTQ files for both cDNA and oligo libraries were processed with DNBelab C Series™ HT Single-Cell Analysis Software (dnbc4tools) to generate raw and filtered gene-by-cell matrices. The Ensembl BDGP 6.46 *Drosophila melanogaster* genome assembly and annotation were used as the reference. dnbc4tools was run with –expectcells 10000 parameter, based on the sequencing report.

#### Quality control and normalization

Gene-by-cell matrices were processed in Seurat (v.5.3.0) (Stuart et al., 2019). Mitochondrial and ribosomal genes were removed because of their strong overrepresentation in the dataset. Ambient RNA contamination was corrected with SoupX (Young and Behjati, 2020). Because Mef2>E(z) KD samples showed higher UMI counts than Mef2>GFP control samples, they were downsampled using scuttle (v1.18.0) to match the controls (McCarthy et al., 2017).

Cells with fewer than 200 and more than 3000 detected features, or more than 7000 UMIs were filtered out.

Downstream processing followed a standard Seurat workflow. Data was normalized and scaled using SCTransform, followed by principal component analysis using 50 principal components. A k-nearest-neighbor graph was constructed with FindNeighbors with dimensions set to 1:35 and k.param = 15. Clustering was performed with FindClusters using the Louvain algorithm with multilevel refinement (resolution = 0.5), and UMAP dimensionality reduction was performed with RunUMAP using dimensions 1:35; number of neighbors set to 15; minimal distance = 0.5 and spread = 1.0.

#### Cluster annotation and validation

Cell clusters were annotated using custom anatomic terms made from BDGP *in situ* gene annotation for embryonic stages 11-12 and 13-16 (https://insitu.fruitfly.org/insitu-mysql-dump/insitu_annot.csv.gz). Enrichment analysis of anatomical terms was performed using the ClusterProfiler (v4.16.0) (Yu et al., 2012) and ontologyIndex (v2.12) (Greene et al., 2017) R packages. To validate cluster annotation, we integrated our dataset with a subset of the snRNA-seq dataset from (Calderon et al., 2022), corresponding to 8-18 h AEL using Harmony (v1.2.3) (Korsunsky et al., 2019) and cluster labels were transferred from the reference dataset using FindTransferAnchors and TransferData from Harmony package in R.

#### Removal of artefactual cell populations

After initial processing, additional filtering steps were taken to remove cell populations lacking proper annotation and suspected to be technical artefacts. First, we identified a large central cluster that could not be assigned to any known cell type, and where marker genes were strongly enriched for ribosomal protein genes. We considered this cluster to represent lower quality cells, and it was therefore removed from both control and Mef2>E(z) KD samples. After this filtering step, 12,970 cells remained in the control sample (median UMI = 560) and 10,133 cells remained in the Mef2>E(z) KD (median UMI = 479).

Second, two clusters enriched with GO terms related to germ line and gonads were overrepresented in the E(z) KD samples. However, Vasa immunostaining did not support an increase in germ cells in the Mef2>E(z) KD samples. These clusters showed enrichment for the S/G2 and M-phase signatures, but staining for phosphorylated H3S10 did not support an increase in mitotic cells in KD embryos. These clusters were not supported by the orthogonal method and therefore excluded from subsequent analyses. After this additional filtering step, 11,461 cells remained in the control sample (median UMI = 574) and 7032 cells remained in the Mef2>E(z) KD sample (median UMI = 509).

#### Marker gene and differential expression analysis

Cluster marker genes were identified using Seurat’s FindAllMarkers function. Differential expression analysis was performed using MAST with the default parameters (v1.33.0) (Finak et al., 2015).

### Immunostaining

For whole-mount immunostaining, embryos were collected on apple juice agar plates supplemented with yeast for 2 hours and aged an additional 14 hours. The embryos were washed, dechorionated in bleach and fixed in formaldehyde as previously described (Haecker et al., 2007). Immunostaining of embryos was performed in PBS containing 0.3% Triton X-100 and 0.5% BSA using rabbit anti-H3K27me3 (Millipore 07-449, 1:100), rabbit anti-Mef2 (1:500, DSHB), rabbit anti-Srp (1:100, kind gift of Ylva Engström), guinea pig anti-Kr (1:300, lab stock). Secondary antibodies used were Alexa Fluor 568-conjugated donkey anti-rabbit (A10042, Thermo Fisher Scientific), Cy3-conjugated donkey anti-guinea pig (706165148, Jackson ImmunoResearch Labs). Embryos were mounted in Vectashield with DAPI (Vector, H-12000). Images were acquired with a Zeiss LSM 800 confocal microscope and single optical sections are shown for all stainings. Quantification of H3K27me3 immunostaining was done on single confocal sections acquired under identical settings of *yw; Mef2-GAL4/UAS-GFP.nls; UAS-E(z) RNAi* and control *yw; Mef2-GAL4/UAS-GFP.nls* 14-16 h AEL embryos in ImageJ. We normalized H3K27me3 mean nuclear signal in muscle nuclei (n = 16-57 GFP-labelled nuclei per section) on mean signal in epidermal nuclei (n= 27-56 nuclei per section).

### RNA in situ hybridization

RNA in situ hybridization was performed on *yw; Mef2-GAL4* and *Mef2-GAL4; UAS-E(z) RNAi* 14-16 h AEL embryos using digoxigenin-labelled antisense RNA probes against *Kr*. RNA in situ hybridization was performed as previously described (Haecker et al., 2007). Embryos were imaged on a Leica DMLB 100T microscope using differential interference contrast with Leica DMC2900 camera.

Images of RNA in situ hybridization for *nau* in wild-type embryos were obtained from the BDGP database (Tomancak et al., 2002).

## Acknowledgements

We thank Ylva Engström, Stockholm University for reagents, Qiaolin Deng and Paulo Jannig for help with MGI sequencing. The authors acknowledge support from the Imaging Facility at Stockholm University (IFSU) and the National Genomics Infrastructure in Stockholm funded by Science for Life Laboratory, the Knut and Alice Wallenberg Foundation and the Swedish Research Council, and SNIC/Uppsala Multidisciplinary Center for Advanced Computational Science for assistance with massively parallel sequencing and access to the UPPMAX computational infrastructure. The computations were enabled by resources provided by the National Academic Infrastructure for Supercomputing in Sweden (NAISS), partially funded by the Swedish Research Council through grant agreement no. 2022-06725.

## Author contributions

SP and MM designed the experiments. SP, AI and JRBW performed the experiments and AP, SP, and AI performed data analysis. SP and MM wrote the original draft with input from all authors. MB and MM acquired funding.

