## Supplemental Figures for "Single-cell chromatin landscapes visualize epigenetic barriers and reveal lineage-specific Polycomb-mediated repression"

**Pirogov et al.**

**Supplementary Figures**

# S1

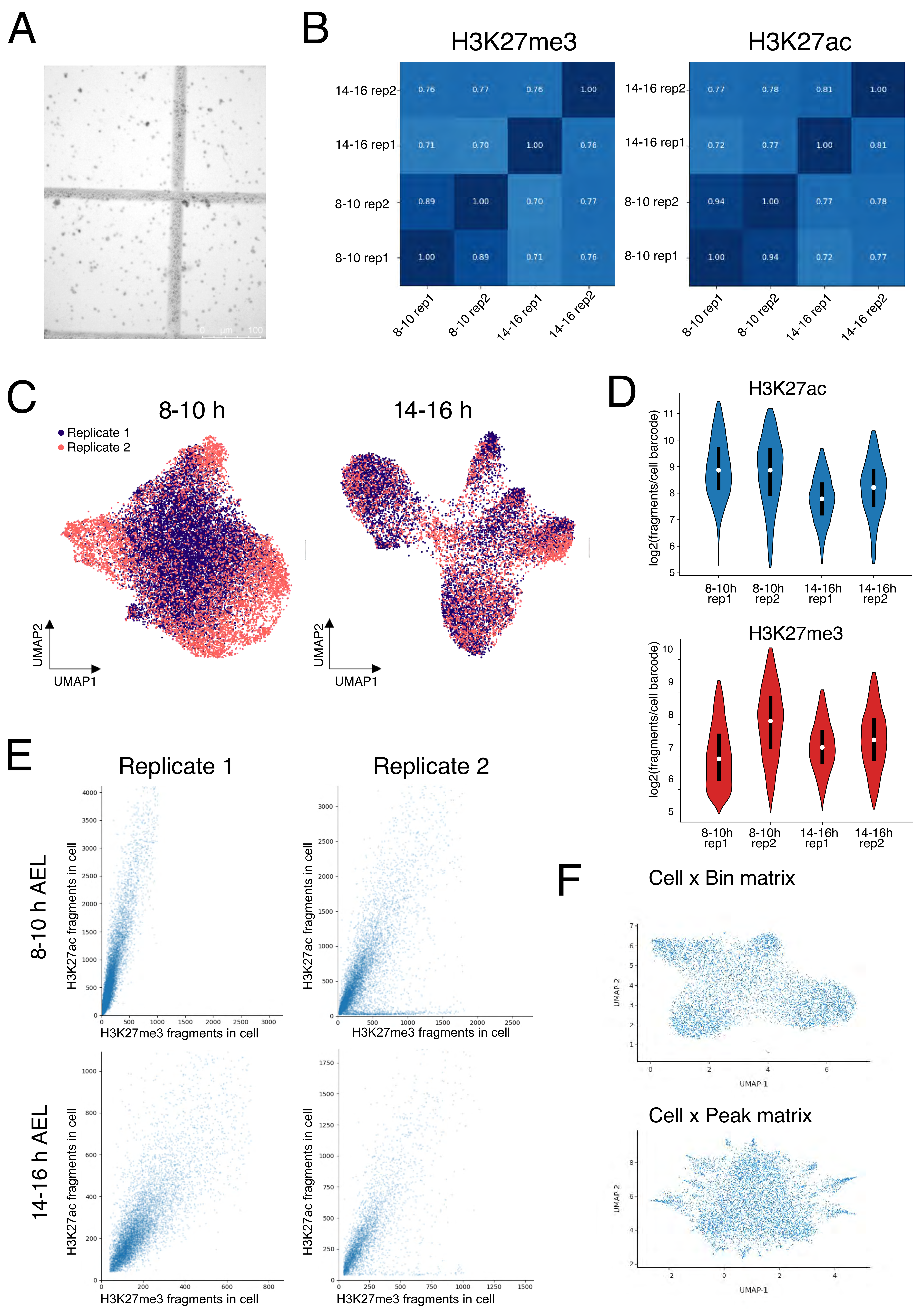
A
B
H3K27me3
H3K27ac
14-16 rep2
14-16 rep1
8-10 rep2
8-10 rep1
8-10 rep1
8-10 rep2
14-16 rep1
14-16 rep2
C
8-10 h
14-16 h
Replicate 1
Replicate 2
UMAP2
UMAP1
D
H3K27ac
H3K27me3
log2(fragments/cell barcode)
8-10h rep1
8-10h rep2
14-16h rep1
14-16h rep2
E
Replicate 1
Replicate 2
8-10 h AEL
14-16 h AEL
H3K27ac fragments in cell
H3K27me3 fragments in cell
F
Cell x Bin matrix
Cell x Peak matrix
UMAP-2
UMAP-1

**Supplementary Figure 1. NanoCUT&Tag libraries are well correlated.**

**A)** An image of the extracted *Drosophila* 14-16 h embryo nuclei stained by Trypan Blue (bright-field). **B)** Pearson correlations between signals (number of normalized fragment counts) in 5kb-bins of two stages in two biological replicates; (left) for H3K27ac, (right) for H3K27me3. **C)** UMAP visualization of bimodal cell-bin matrix of nanoCT nuclei colored by replicate and split by stage. **D)** Violin plots of unique fragments per nucleus for both stages in two biological replicates; (top) fragments with the H3K27ac modality barcode, (bottom) fragments with H3K27me3 modality barcode. **E)** Scatter plot of number of H3K27ac and H3K27me3 fragments per each cell barcode. **F)** Comparison between UMAP visualizations of bimodal cell-genomic bin and cell-peak matrices for H3K27ac modality stage 14-16h, replicate 1.

S2

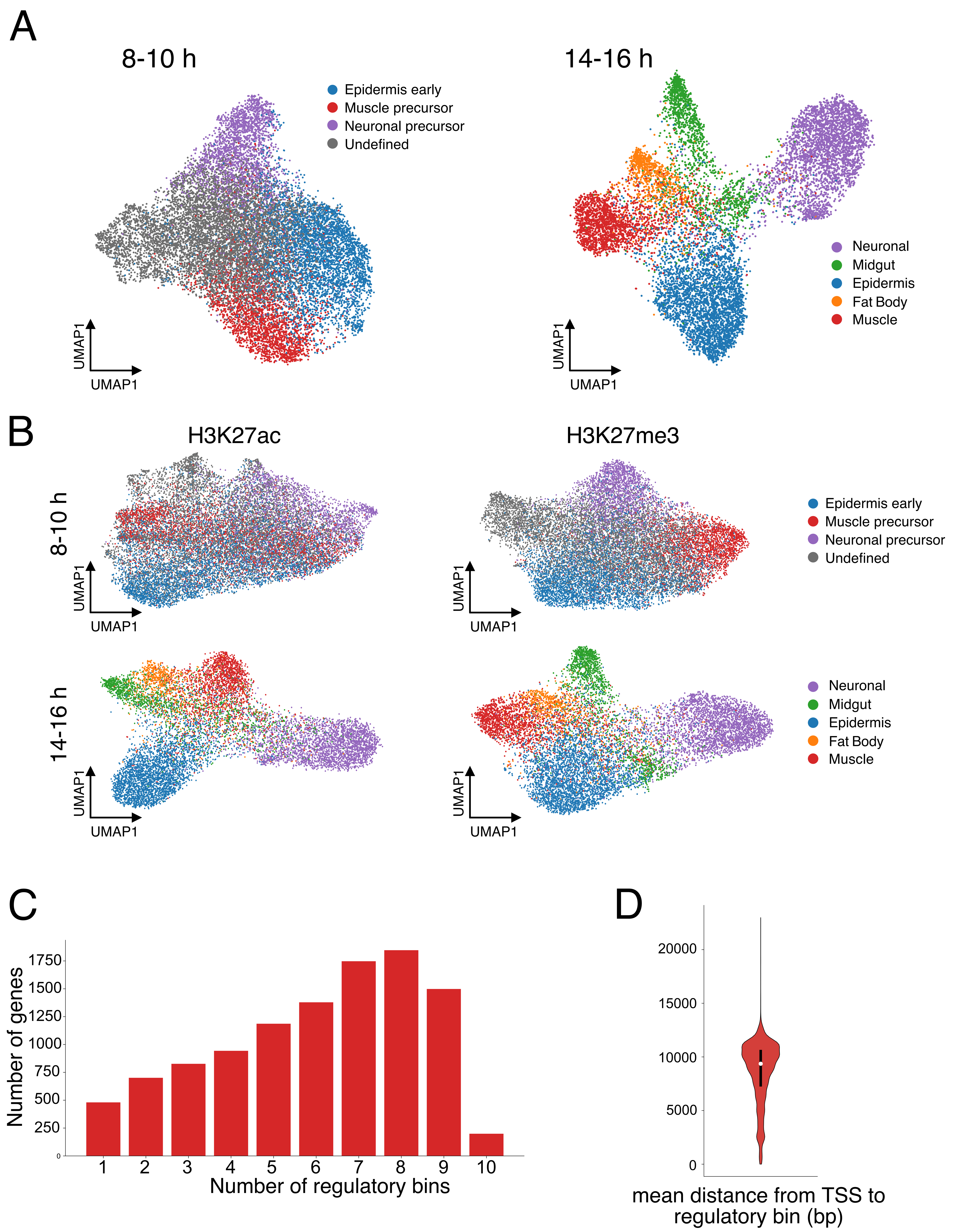
A
8-10 h
Epidermis early
Muscle precursor
Neuronal precursor
Undefined
UMAP1
UMAP1
14-16 h
Neuronal
Midgut
Epidermis
Fat Body
Muscle
UMAP1
UMAP1
B
H3K27ac
H3K27me3
8-10 h
14-16 h
Epidermis early
Muscle precursor
Neuronal precursor
Undefined
Neuronal
Midgut
Epidermis
Fat Body
Muscle
UMAP1
UMAP1
C
Number of genes
1750
1500
1250
1000
750
500
250
0
1
2
3
4
5
6
7
8
9
10
Number of regulatory bins
D
20000
15000
10000
5000
0
mean distance from TSS to regulatory bin (bp)

**Supplementary Figure 2. Bimodal nanoCUT&Tag epigenomic profiling outperforms unimodal profiling.**

**A)** UMAP visualization of bimodal cell-bin matrix of nanoCT nuclei colored by annotated cell cluster and split by stage. **B)** UMAP visualizations of stage-specific and modality-specific embeddings of unimodal cell-bin matrix of nanoCT nuclei colored by Leiden clusters from S2A. **C)** Histogram of number of 5kb-bins connected to genes after scRNA-seq inference (the bin overlapping a promoter of any gene is included by default). **D)** Violin plot of distance distribution between TSS and connected bins.

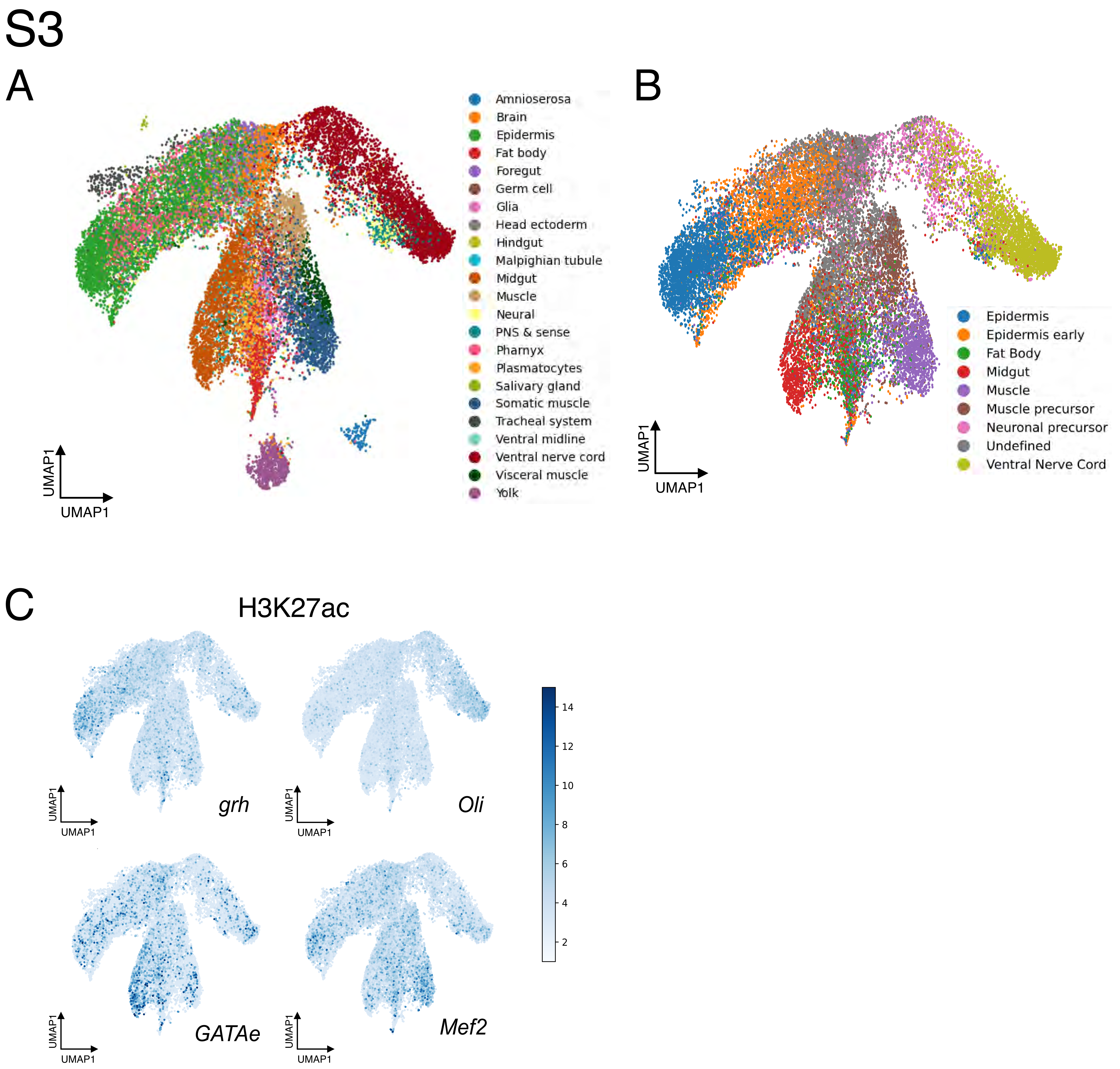
S3
A
Amnioserosa
Brain
Epidermis
Fat body
Foregut
Germ cell
Glia
Head ectoderm
Hindgut
Malpighian tubule
Midgut
Muscle
Neural
PNS & sense
Pharnyx
Plasmatocytes
Salivary gland
Somatic muscle
Tracheal system
Ventral midline
Ventral nerve cord
Visceral muscle
Yolk
UMAP1
UMAP1
B
Epidermis
Epidermis early
Fat Body
Midgut
Muscle
Muscle precursor
Neuronal precursor
Undefined
Ventral Nerve Cord
UMAP1
UMAP1
C
H3K27ac
grh
Oli
GATAe
Mef2
14
12
10
8
6
4
2

**Supplementary Figure 3. Integration of nanoCUT&Tag with scATAC-seq.**

**A)** Label transfer of scATAC-seq clusters to the scATAC-seq nuclei integrated with nanoCT nuclei. **B)** Label transfer of nanoCT clusters (from Fig.1C) to the nanoCT nuclei integrated with scATAC-seq nuclei. **C)** UMAP visualization of cell-bin matrix of nanoCT nuclei after integration, coloured by H3K27ac level at connected genomic bins, for a subset of cell-type specific transcription factors.

S4

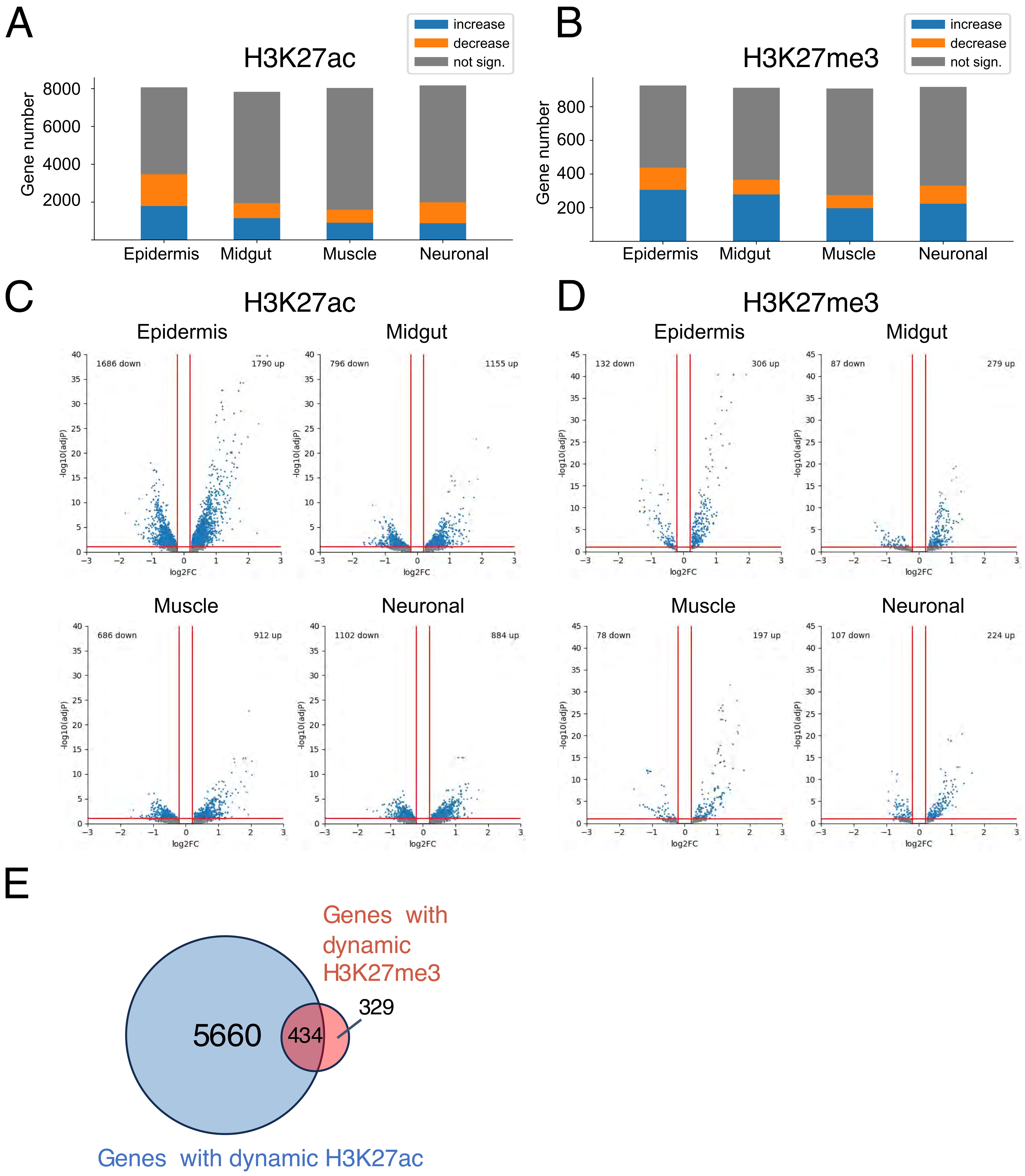

A
H3K27ac
increase
decrease
not sign.
Gene number
8000
6000
4000
2000
Epidermis
Midgut
Muscle
Neuronal
B
H3K27me3
increase
decrease
not sign.
Gene number
800
600
400
200
Epidermis
Midgut
Muscle
Neuronal
C
H3K27ac
Epidermis
1686 down
1790 up
Midgut
796 down
1155 up
Muscle
686 down
912 up
Neuronal
1102 down
884 up
-log10(adjP)
log2FC
D
H3K27me3
Epidermis
132 down
306 up
Midgut
87 down
279 up
Muscle
78 down
197 up
Neuronal
107 down
224 up
-log10(adjP)
log2FC
E
Genes with dynamic H3K27me3
5660
434
329
Genes with dynamic H3K27ac

**Supplementary Figure 4. Changes to H3K27ac and H3K27me3 over developmental time.**

**A,B)** Barplots of genes with **A)** H3K27ac signal or **B)** H3K27me3 signal split by increasing, decreasing , and no significant change of signal for each mark between 8-10 h and 14-16 h old embryos ($|\log_2 FC| > 0.2$, p-value $< 0.1$). **C,D)** Volcano-plots showing genes with significant change in **C)** H3K27ac or **D)** H3K27me3 signal. **E)** Venn diagram of genes significantly changing in H3K27ac, H3K27me3 or both.

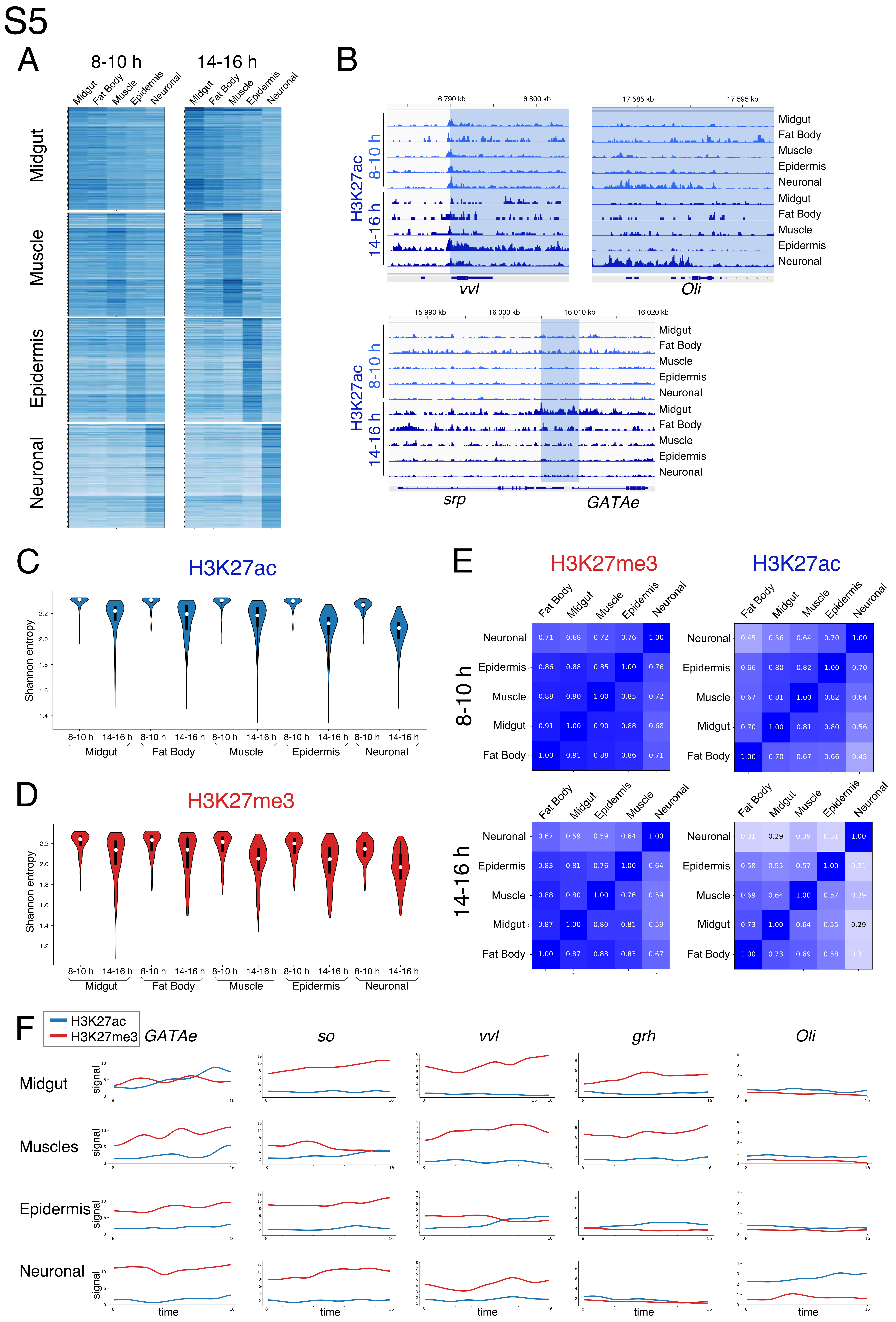
S5
A
8-10 h
14-16 h
Midgut
Fat Body
Muscle
Epidermis
Neuronal
B
H3K27ac 8-10 h
H3K27ac 14-16 h
6 790 kb
6 800 kb
17 585 kb
17 595 kb
vvl
Oli
15 990 kb
16 000 kb
16 010 kb
16 020 kb
srp
GATAe
C
H3K27ac
Shannon entropy
D
H3K27me3
E
H3K27me3
H3K27ac
8-10 h
14-16 h
F
H3K27ac
H3K27me3
GATAe
so
vvl
grh
Oli
Midgut
Muscles
Epidermis
Neuronal
signal
time

**Supplementary Figure 5. Increased cell-type specificity of H3K27ac during development.**

**A)** Heatmap of normalized H3K27ac counts at connected bins of top 200 cell-type specifically acetylated genes in two embryo stages, across cell clusters. **B)** Genome browser snapshots showing pseudobulk H3K27ac signal in different cell-types for the *vvl, Oli, srp* and *GATAe* genes in two embryo stages. **C, D)** Violin plots showing Shannon entropy of H3K27ac (**C**) and H3K27me3 (**D**) in the top 200 cell-type-specifically H3K27-acetylated or -methylated genes. **E)** Pearson correlation between (left) H3K27me3 or (right) H3K27ac signals in different cell types at bins connected to genes in two embryo stages. Only genes with a signal above the threshold are included (8-10h H3K27ac total genes = 11726; 14-16h H3K27ac total genes = 11636; 8-10h H3K27me3 total genes = 772; 14-16h H3K27me3 total genes = 709). **F)** Plots showing the smoothed H3K27ac and H3K27me3 signals for representative genes expressed in midgut (*GATAe*), muscle (*Mef2*), epidermis (*vvl, grh*) and neurons (*Oli*) along inferred developmental time.

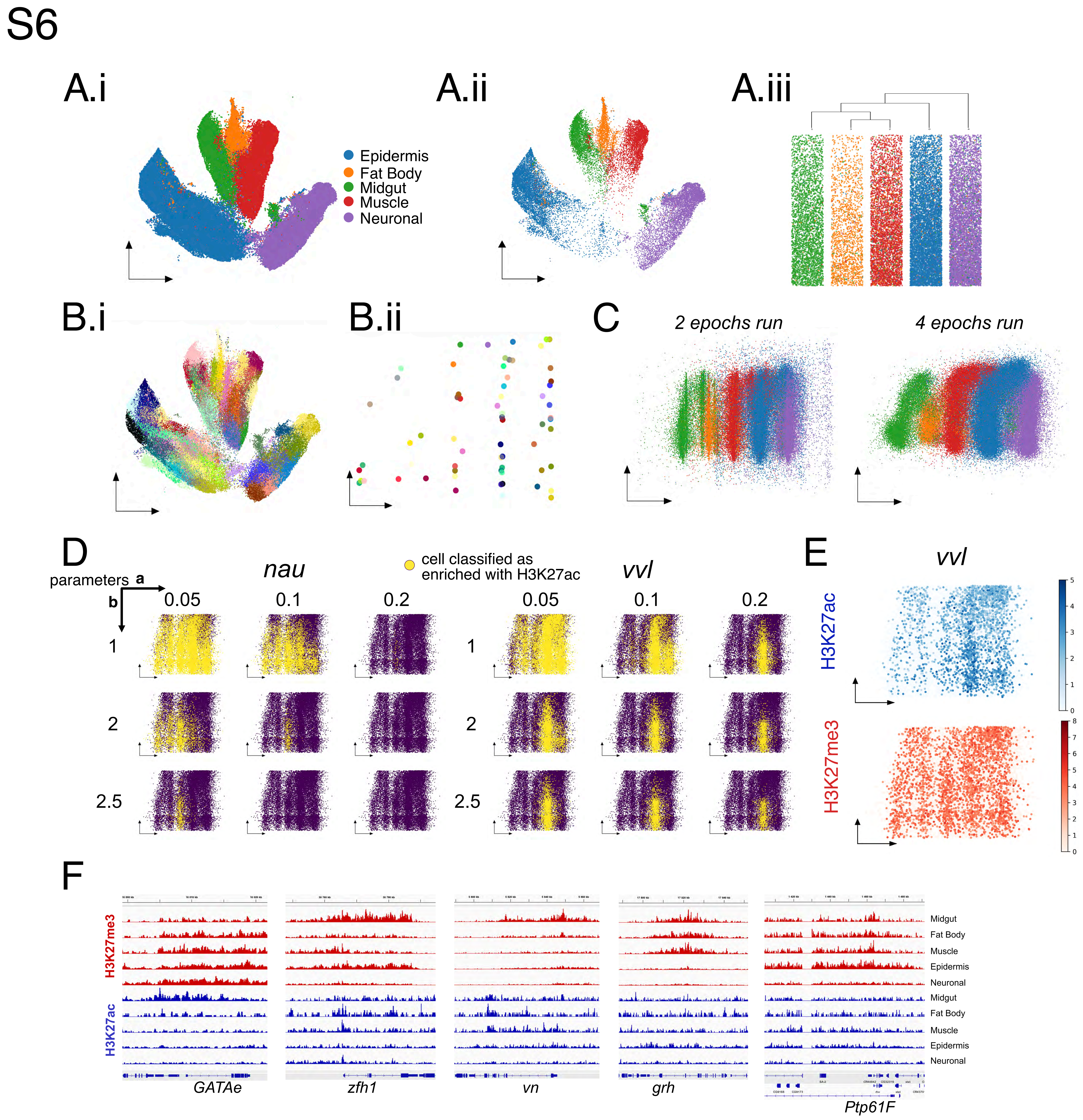
S6
A.i
A.ii
A.iii
Epidermis
Fat Body
Midgut
Muscle
Neuronal
B.i
B.ii
C
2 epochs run
4 epochs run
D
parameters a
b
nau
vvl
cell classified as enriched with H3K27ac
0.05
0.1
0.2
1
2
2.5
E
H3K27ac
H3K27me3
F
Midgut
Fat Body
Muscle
Epidermis
Neuronal
GATAe
zfh1
vn
grh
Ptp61F

**Supplementary Figure 6. Generation of gene-specific epigenetic landscapes.**

**Ai)** UMAP visualization of cell-genomic bin matrix of scATAC-seq nuclei from 8-16 hours embryos after integration with nanoCT nuclei, colored by cell type. **Aii)** Only late stage (14-16 h) cells are shown, representing the terminal states. **Aiii)** Dendrogram of hierarchically ordered clusters. **Bi)** UMAP visualization as in **Ai** colored by metacells. **Bii)** Metacells on the cell-time embedding, placed according to their similarity to the terminal clusters and mean cell age. **C)** Cell type-developmental time embedding of scATAC-seq nuclei placed according to their individual predicted cell ages on y (time)-axis, and according to their respective metacell position along x-axis, with different numbers of UMAP iterations (N of epochs = 2,4). **D)** Cell-time embeddings demonstrating the result of classification for H3K27ac enrichment at bins connected to *nau* and *vvl* depending on a and b coefficients (see Methods). **E)** Cell-time embedding for the *vvl* gene with H3K27ac (top) and H3K27me3 (bottom) signal used for classification. **F)** Genome browser snapshots with pseudobulk H3K27me3 and H3K27ac signals in 14-16 h embryos for the respective genes.

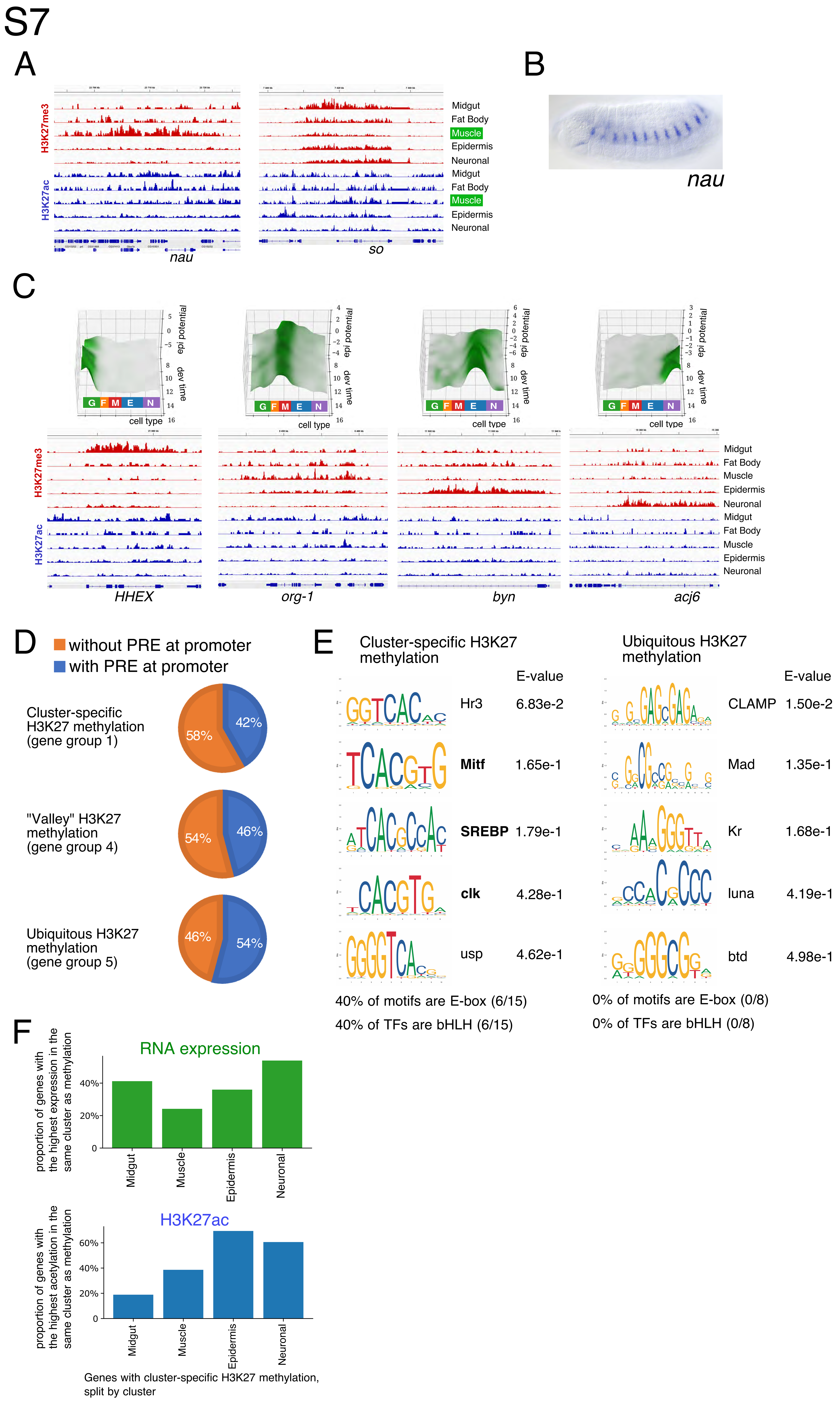

S7
A
H3K27me3
H3K27ac
Midgut
Fat Body
Muscle
Epidermis
Neuronal
Midgut
Fat Body
Muscle
Epidermis
Neuronal
nau
so
B
nau
C
epi potential
dev time
cell type
G F M E N
H3K27me3
H3K27ac
Midgut
Fat Body
Muscle
Epidermis
Neuronal
Midgut
Fat Body
Muscle
Epidermis
Neuronal
HHEX
org-1
byn
acj6
D
without PRE at promoter
with PRE at promoter
Cluster-specific H3K27 methylation (gene group 1)
58%
42%
"Valley" H3K27 methylation (gene group 4)
54%
46%
Ubiquitous H3K27 methylation (gene group 5)
46%
54%
E
Cluster-specific H3K27 methylation
E-value
Hr3 6.83e-2
Mitf 1.65e-1
SREBP 1.79e-1
clk 4.28e-1
usp 4.62e-1
40% of motifs are E-box (6/15)
40% of TFs are bHLH (6/15)
Ubiquitous H3K27 methylation
E-value
CLAMP 1.50e-2
Mad 1.35e-1
Kr 1.68e-1
luna 4.19e-1
btd 4.98e-1
0% of motifs are E-box (0/8)
0% of TFs are bHLH (0/8)
F
RNA expression
proportion of genes with the highest expression in the same cluster as methylation
40%
20%
0
Midgut
Muscle
Epidermis
Neuronal
H3K27ac
proportion of genes with the highest acetylation in the same cluster as methylation
60%
40%
20%
0
Midgut
Muscle
Epidermis
Neuronal
Genes with cluster-specific H3K27 methylation, split by cluster

**Supplementary Figure 7. Co-occurrence of H3K27me3, H3K27ac and RNA expression.**

**A)** Genome browser snapshots with pseudobulk H3K27me3 and H3K27ac signals in 14-16 h embryos at the *nau* and *so* loci. **B)** *nau* in situ hybridization staining image from BGDP (Tomancak et al., 2002). **C)** 3D epigenetic landscapes and genome browser shots with pseudobulk H3K27me3 and H3K27ac signals in 14-16 h embryos for genes *HHEX* expressed in midgut, *org-1* (muscle), *byn* (epidermis) and *acj6* (neurons). **D)** Pie charts of genes with or without a PRE in proximity of their promoters (1kb around TSS) for genes in group 1 (cluster-specific H3K27 methylation), group 4 (valley pattern), and group 5 (ubiquitously H3K27 methylated). **E)** Transcription factor motifs enriched in promoter regions (1kb around TSS) of genes in group 1 and group 5. bHLH TFs are highlighted in bold. **F)** Bar plot of the ratio of genes with highest gene expression (top) or highest H3K27 acetylation (bottom) in the same cell cluster as genes with cluster-specific H3K27 methylation (group 1).

# S8

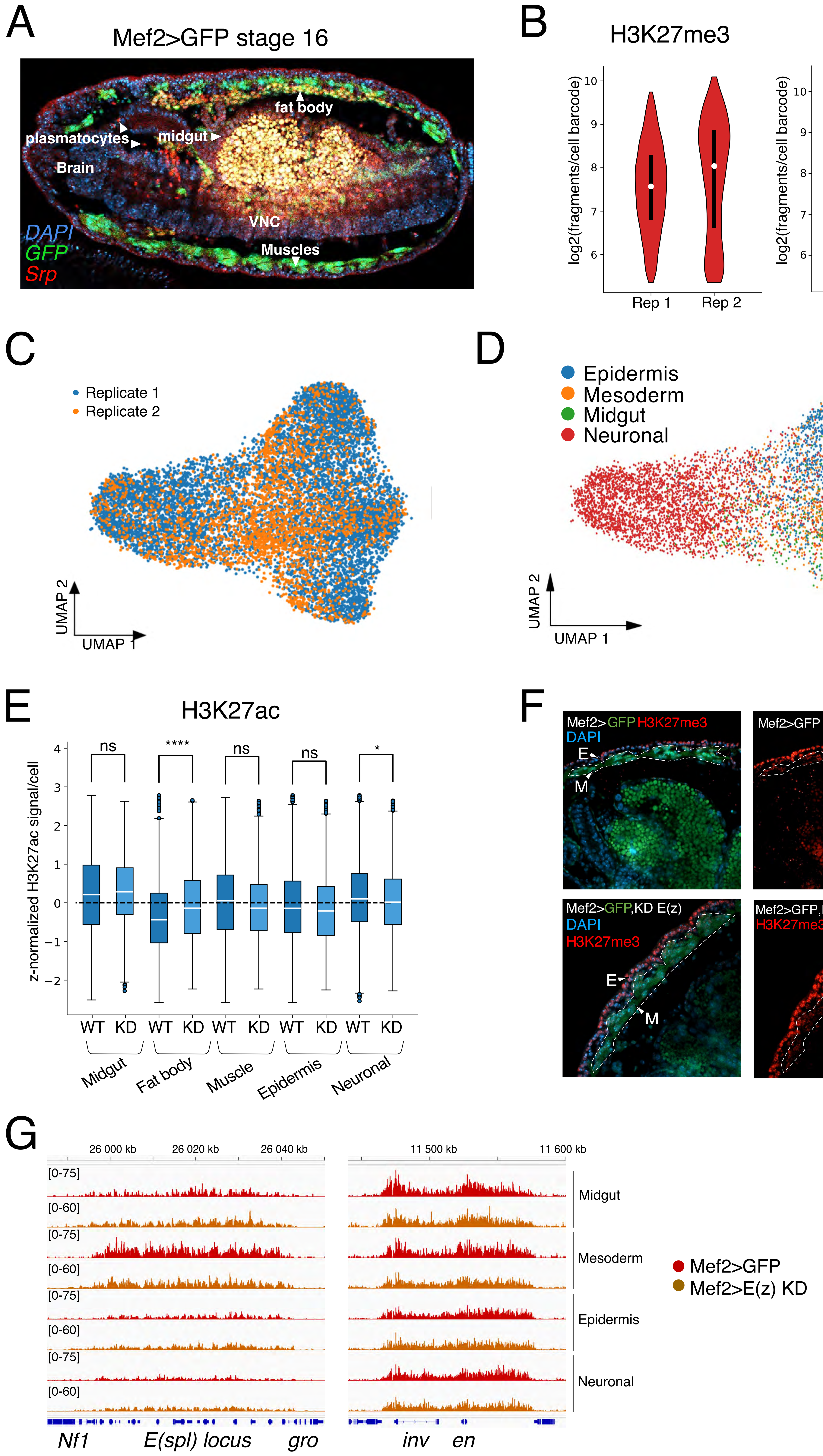
A
Mef2>GFP stage 16
fat body
plasmatocytes
midgut
Brain
VNC
Muscles
DAPI
GFP
Srp
B
H3K27me3
H3K27ac
log2(fragments/cell barcode)
Rep 1
Rep 2
C
Replicate 1
Replicate 2
UMAP 2
UMAP 1
D
Epidermis
Mesoderm
Midgut
Neuronal
E
H3K27ac
z-normalized H3K27ac signal/cell
ns
****
ns
ns
*
WT
KD
Midgut
Fat body
Muscle
Epidermis
Neuronal
F
Mef2>GFP H3K27me3
DAPI
E
M
Mef2>GFP,KD E(z)
H3K27me3
G
26 000 kb
26 020 kb
26 040 kb
11 500 kb
11 600 kb
[0-75]
[0-60]
Midgut
Mesoderm
Epidermis
Neuronal
Mef2>GFP
Mef2>E(z) KD
Nf1
E(spl) locus
gro
inv
en

**Supplementary Figure 8. Muscle-specific knock-down of the H3K27 methyltransferase E(z).**

**A)** Confocal cross-section of stage 16 Mef2>GFP embryo stained with DAPI (blue) and anti-Srp (fat body marker in red). GFP is in green. Ventro-lateral view of the embryo with tissues indicated. **B)** Violin plots of unique fragments per nucleus for 14-16 h Mef2>E(z)KD embryos in two biological replicates; (left) fragments with the H3K27ac modality barcode, (right) fragments with H3K27me3 modality barcode. **C)** UMAP visualization of bimodal cell-genomic bin matrix of nanoCT stage 16 Mef2>E(z)KD nuclei colored by biological replicate. **D)** UMAP visualization of nanoCT nuclei colored by annotated cell clusters without integration with wild-type embryos. **E)** Boxplots of z-normalized H3K27ac signal per cell across all bins, comparing wild-type (WT) with Mef2>E(z)KD within each cell type. * indicates p-value ≤ 0.05 and **** indicates p-value ≤ 0.0001, and ns stands for not significant (two-tailed Student's t-test with Benjamini-Hochberg correction) **F)** Confocal sections from stage 16 Mef2>GFP control and Mef2>E(z)KD embryos stained for H3K27me3. Top: control; bottom: E(z)KD in muscle. Epidermis (E) and muscle (M) are indicated. **G)** Genome browser shots of pseudobulk H3K27me3 signal in wild type and Mef2>E(z) KD embryos at two loci (*E(spl)* and *inv-en*).

S9

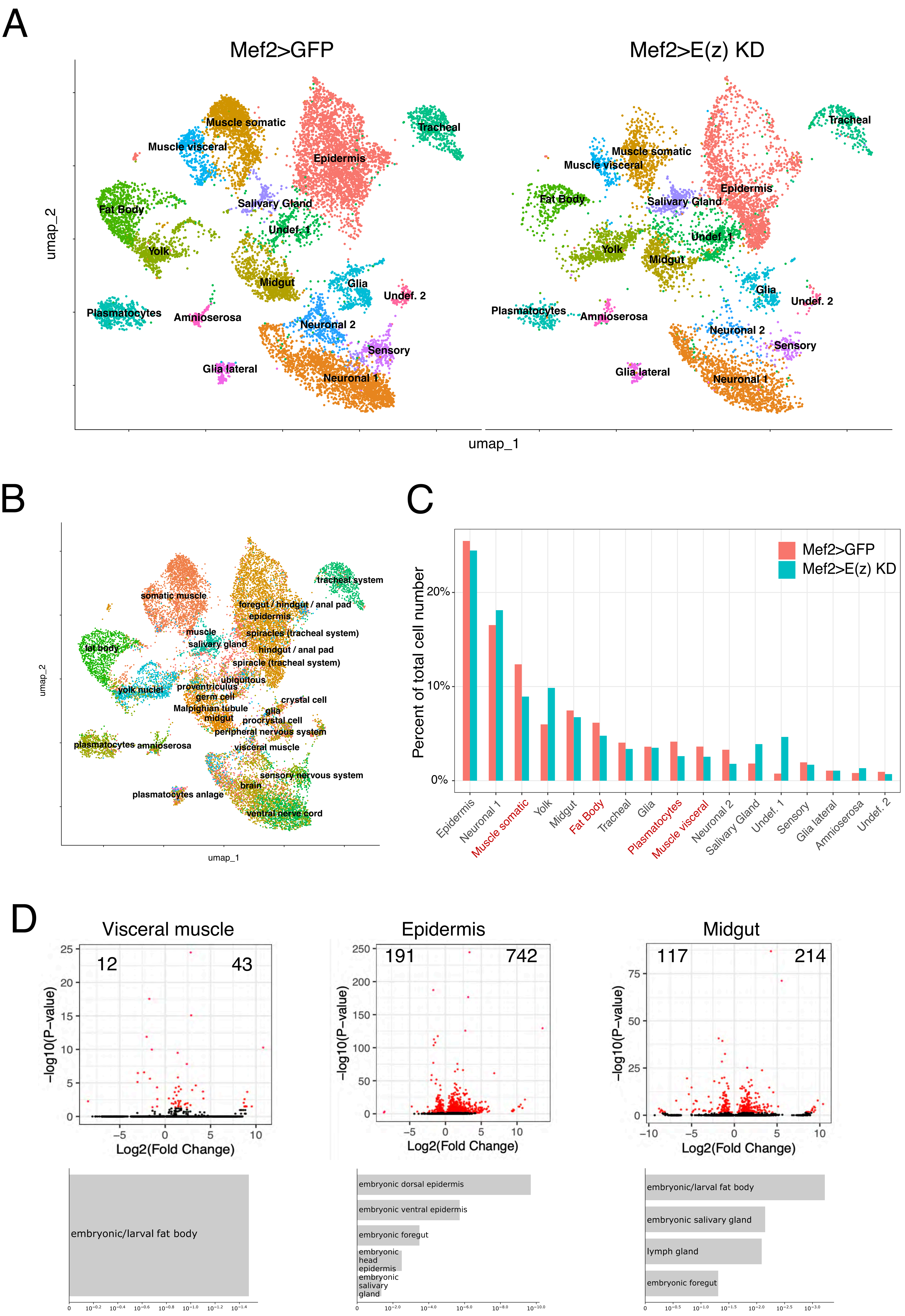

A
Mef2>GFP
Mef2>E(z) KD
umap_2
umap_1
Muscle somatic
Muscle visceral
Tracheal
Epidermis
Fat Body
Salivary Gland
Undef. 1
Yolk
Midgut
Glia
Undef. 2
Plasmatocytes
Amnioserosa
Neuronal 2
Sensory
Glia lateral
Neuronal 1
B
somatic muscle
tracheal system
foregut / hindgut / anal pad
epidermis
muscle
spiracles (tracheal system)
salivary gland
hindgut / anal pad
fat body
spiracle (tracheal system)
ubiquitous
yolk nuclei
proventriculus
germ cell
crystal cell
Malpighian tubule
glia
midgut
procrystal cell
peripheral nervous system
plasmatocytes
amnioserosa
visceral muscle
sensory nervous system
brain
plasmatocytes anlage
ventral nerve cord
C
Percent of total cell number
Mef2>GFP
Mef2>E(z) KD
20%
10%
0%
Epidermis
Neuronal 1
Muscle somatic
Yolk
Midgut
Fat Body
Tracheal
Glia
Plasmatocytes
Muscle visceral
Neuronal 2
Salivary Gland
Undef. 1
Sensory
Glia lateral
Amnioserosa
Undef. 2
D
Visceral muscle
12
43
−log10(P−value)
Log2(Fold Change)
Epidermis
191
742
Midgut
117
214
embryonic/larval fat body
embryonic dorsal epidermis
embryonic ventral epidermis
embryonic foregut
embryonic head epidermis
embryonic salivary gland
embryonic salivary gland
lymph gland

**Supplementary Figure 9. Single-nucleus RNA-seq of control and Mef2>E(z) knockdown embryos.**

**A)** UMAP visualization of snRNA-seq split by condition, colored by cell-type annotation (two biological replicates merged). **B)** Label transfer from 14-16 h scRNA-seq (Calderon et al., 2022) to our snRNA-seq after integration. **C)** Bar plot of cell type fractions in Mef2>GFP control and Mef2>E(z)KD embryos (two biological replicates merged). Mesodermal lineages are highlighted in red. **D)** Volcano plot for differential gene expression between Mef2>GFP control and Mef2>E(z) KD embryos in the visceral muscle, epidermis, and midgut clusters. Enriched anatomic terms in up-regulated genes are shown below each cluster.

# S10

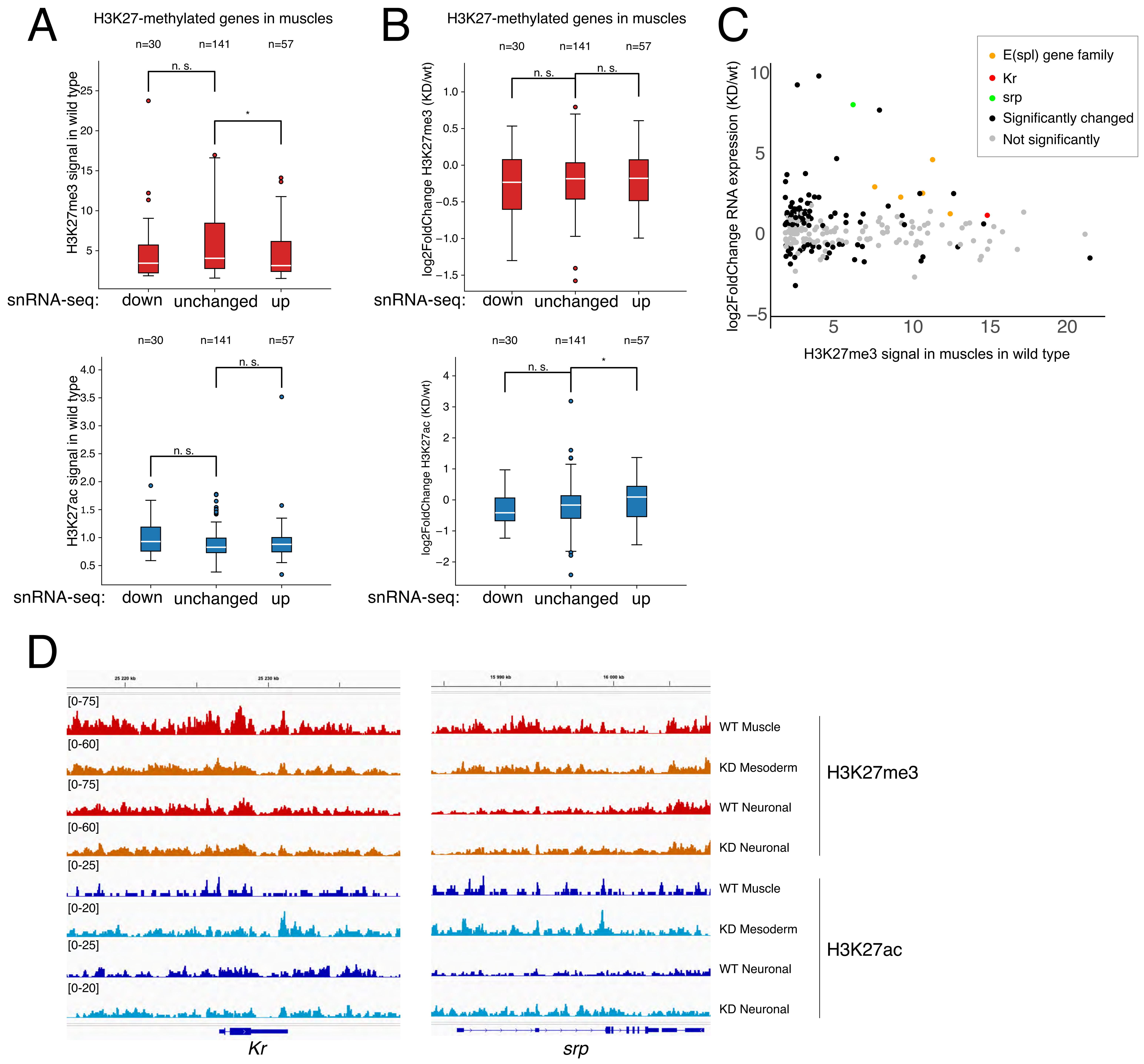
A
H3K27-methylated genes in muscles
n=30
n=141
n=57
n. s.
*
H3K27me3 signal in wild type
snRNA-seq: down unchanged up
n=30
n=141
n=57
n. s.
n. s.
H3K27ac signal in wild type
snRNA-seq: down unchanged up
B
H3K27-methylated genes in muscles
n=30
n=141
n=57
n. s.
n. s.
log2FoldChange H3K27me3 (KD/wt)
snRNA-seq: down unchanged up
n=30
n=141
n=57
n. s.
*
log2FoldChange H3K27ac (KD/wt)
snRNA-seq: down unchanged up
C
E(spl) gene family
Kr
srp
Significantly changed
Not significantly
log2FoldChange RNA expression (KD/wt)
H3K27me3 signal in muscles in wild type
D
[0-75]
[0-60]
[0-75]
[0-60]
[0-25]
[0-20]
[0-25]
[0-20]
WT Muscle
KD Mesoderm
WT Neuronal
KD Neuronal
H3K27me3
WT Muscle
KD Mesoderm
WT Neuronal
KD Neuronal
H3K27ac
Kr
srp

**Supplementary Figure 10. Gene expression changes in muscle from Mef2>E(z) knockdown embryos.**

**A)** Boxplots of H3K27me3 and H3K27ac signal at H3K27me3 decorated genes in muscle in wild-type 14-16 h embryo for up-regulated, down-regulated, or not-significantly changed genes (based on snRNA-seq). * indicates p-value ≤ 0.05, ns indicates no significant change. **B)** Boxplots of $\log_2$ fold change for H3K27ac and H3K27me3 signal between Mef2>E(z)KD embryos and wild type embryos in muscle for the same group of genes as in (**A**). **C)** MA-plot of H3K27me3 signal in wild-type 14-16 h embryonic muscle and $\log_2$ fold change of RNA expression in somatic muscles. Genes encoding the transcription factors *Kr*, *srp*, and *E(spl)* are highlighted. **D)** Genome browser screenshots of H3K27me3 and H3K27ac at *Kr* and *srp* loci in wild-type (WT) and Mef2>E(z)KD embryos.
