## Extended Methods for "Single-cell chromatin landscapes visualize epigenetic barriers and reveal lineage-specific Polycomb-mediated repression"

### 1 NanoC&T data processing

We create a fragment matrix by assigning all fragments into 5000 bp bins. For embedding our data, we used the multiview spectral algorithm from SnapATAC2, which combines information from both modalities. We remove one (almost always first) principal component (PC) where the PC correlates $|\rho| > 0.6$ with meth/acet summed counts. We reasoned that this PC likely contains mostly information that is directly related to fragment count per cell.
We annotated cells using gene in-situ annotations database (BDGP). The BDGP data was formatted as a custom GO Term namespace (called anatomic terms). We then ran `github:tanghaibao/goatools` (GOATOOLS) on this custom namespace to get GO Term statistics on the in-situ data. In this way, we could create lists of enriched anatomic terms per cell cluster to aid annotation.

#### 1.1 Regulatory bin-to-gene connection.

We ran `github:gao-lab/GLUE` to integrate our NanoC&T data with ATAC or RNA data.
To integrate RNA-seq with ATAC-seq, we created a graph connecting promoter to ±20 kb.

$$w = (\frac{|\text{promoter} - \text{bin}| + 500}{500})^{-0.75}, |\text{promoter} - \text{bin}| < 20\,000$$

Then, the cell × gene matrix is created from taking feature matrix $X$,

$$X \in \mathbb{R}^{n \times m}, \text{ where } n, m \text{ are cells, bin features}$$

$$X_{gene} = XRD, \text{ where } R \in \{0,1\}^{m \times k}, \text{ where } k \text{ is gene features; } D = \text{diag}(\frac{1}{\mathbf{1}^T R})$$

### 2 Landscape embedding

We want to create an embedding with two dimensions, namely ”developmental time” and ”cell type”. Developmental time was straightforward, as we had a neural-network inferered age for all our cells after integrating with published ATAC-seq.
For creating the cell type dimension, we wanted it to preserve a global structure, but allow the one dimensional cell type value to be continuous. We experimented first with 1D UMAP, but this introduced the time element into the dimension, which we wanted to avoid. A possible solution would have been to regress out time before embedding, but this would probably have been impossible given our extremely low fragments per cell. Another solution would have been to remove the dimensions correlated with time from the embedding, but this would probably be impractical if the time information is spread between multiple components and we cannot isolate one single component consisting with mostly time information. Our embedding is low-resolution and would probably become even worse after removing any components correlating with developmental time.
To address these problems, we create an algorithm that

1. Determines late cell types (leiden clusters on late subset of data) $L = \{i : t_i > \tau\}$, where $\tau$ is the time threshold.
2. Creates a hierarchical scaffold on these cells (ordered).
3. Makes new metacell clusters on whole dataset.
4. Assings metacells to scaffold based on $k$NN-graph.
5. Expands metacells to single cell resolution.

For step 2, we ordered the leaves of the scaffold dendrogram using the algorithm outlined in `arXiv:2306.00833v2`. It works by

a) computing the similarity between all pairs of bottom clusters;

b) merging the two clusters that are the most similar, and updating the similarities between this new cluster and the existing ones;

c) repeating step (b) until all clusters have been merged into a single one.

In step 4, the x-coordinate for each metacell (cluster) $j$ is computed as:

$$w_{ji} = \sum_{u \in C_j} \sum_{v \in S_i} (A_{uv} + A_{vu})$$

where $A$ is the $k$NN adjacency matrix, $C_j$ cells in cluster $j$, and $S_i$ cells in scaffold $i$.

$$x_j = \frac{\sum_{i=1} x_i \cdot w_{ji}}{\sum_{i=1} w_{ji}},$$

where $x_i$ are the scaffold leaf positions, $x_i = \text{order}_i$.
The y-coordinate for each cluster $j$ is the negative mean of developmental time (or other y-axis variable):

$$y_j = -\bar{t}_j$$

The single-cell resolution landscape is gotten from these cluster coordinates,

$$\mathcal{E}_{init} = \left( \sum_{j=1} x_j \cdot \mathbf{1}_{C_j}, \sum_{j=1} y_j \cdot \mathbf{1}_{C_j} \right)$$

where $\mathbf{1}$ is the indicator function.

$$\mathcal{E} = \text{UMAP}(A|\text{init} = \mathcal{E}_{init}; \text{epochs} = n)$$

where $n$ is a number between 5-10, leading to a slightly more optimized state of embedding, but not one that has converged on UMAP terms.
The algorithm works best when run on a large number of cells, why we used the ATAC data we had integrated with our NanoC&T data for creating the landscape. After creating the embedding, we could simply fit a KNeighborsRegressor (`sklearn`) on the GLUE embedding of ATAC, and after fitting the regressor, predict the landscape for NanoC&T data on the GLUE embedding.

#### 2.1 3D Landscape plot

We want to visualize the methylation level, the acetylation, or an epigenetic potential $f = \text{methylation} - \text{acetylation}$ on this newly created embedding. The easiest way for us to do this was to make a rectangular grid and infer the value at each gridcell by the cells that were inside that gridcell (creating so called griddata). A grid $g_x \times g_y$ is created, spanning the landscape embedding min, max x- and y-coordinates.

$$C \in \mathbb{N}_0^{g_x \times g_y}$$
$$Z \in \mathbb{R}^{g_x \times g_y \times m}$$

where $m$ is the number of features (genes). Matrix $C$ is the number of cells in each gridcell. $Z$ is the mean feature counts over all cells in the gridcell.
To aid visualization, further transformations on $Z$ are done. A median filter followed by a gaussian filter. Both filters are slightly modified to add support for matrices with missing data, which in our case are gridcells with number of cells not reaching the threshold; these were set to 0; however, they should be treated differently from a bona-fide 0, warranting modification of the filter algorithms.
The median filter has an arbitrary window size of $k \times k$ grid cells, and instead of calulating the median at each grid cell, it calculates the median excluding missing values, that is, including only $C_{ij} \geq \text{min\_cells}$. The modified gaussian blur works on a similar principle.

##### 2.1.1 Modified Gaussian Blur

First, we zero out positions where the cell count is below the threshold:

$$Z_{ij} = \begin{cases} 0 & \text{if } C_{ij} < \text{min\_cells} \\ Z_{ij} & \text{otherwise} \end{cases}$$

Apply a Gaussian filter with kernel $G_\sigma$ (bandwidth $\sigma$, 0-padded on all sides):

$$Z = G_\sigma * Z$$

where the convolution is applied over all dimensions.
To correct for the effect introduced by zeroing missing values, we compute normalization weights by convolving the indicator of non-missing positions:

$$N = G_\sigma * (C \geq \text{min\_cells})$$

Finally, impute the data matrix:

$$Z_{ij} = \begin{cases} \text{nan\_fill} & \text{if } N_{ij} < \epsilon \\ \dfrac{Z_{ij}}{N_{ij}} & \text{otherwise} \end{cases}$$

where nan_fill is a value to fill cells that remain 0 even after imputation.

###### 2.1.2 RNA Activity on 3D landscape

To plot RNA activity onto the landscape, we used a custom colormap that took the RNA level and assigned it a green color reflecting its relative expression level. It was blended with gray that was a function of the z-value of the landscape to make the topography easier to see.
However, to show RNA activity, the RNA data had to first be connected with the NanoC&T data. We used a KNeighborsRegressor (`sklearn`), which we fit on the joint embedding of ATAC and RNA to predict the landscape for RNA. After both NanoC&T and RNA data had been predicted a landscape, we could simply overlay them.

### 3 Mark classification

We wanted to classify whether a cell has or has not a mark, looking at a genomic region of interest. We reasoned that we could create an activity score for all cells over all genes, and use a threshold to determine the binary activity of cells in the context of a certain gene. To focus on the manifold of data instead of the particular datapoints, we first create a normalized activity matrix that takes into account cell communities. These communities could have been calculated with Leiden, but Leiden is not random enough for our purposes, favouring stable communities instead of completely random ones. Thus, we used a random cutting algorithm on the $k$NN graph.

#### 3.1 Normalized activity matrix

First, we make a binarized fragment matrix $X_B \in \{0,1\}^{n \times m}$ by picking the top 20% of fragments from normalized matrix $B$:

$$X_B = \begin{cases} 1 & X > Q_{80}(X) \\ 0 & \text{otherwise} \end{cases}$$

The activity matrix $A \in \mathbb{R}^{n \times m}$ is calculated as

$$A = LD(L^T X_B)$$

where $D = \text{diag}(\frac{1}{\mathbf{1}^T L})$
We partition the $k$NN graph $p$ times to get a membership matrix

$$L \in \{0,1\}^{n \times p}$$

###### 3.1.1 Random communities

Let $G = (V, E)$ be a $k$NN graph with $n$ vertices. We wish to partition the set of vertices $V$ into $k$ disjoint clusters $C_1, C_2, ..., C_k$.
The algorithm operates by creating one cluster at a time from the set of currently unassigned nodes.
Let $U_1 = V$ be the set of all nodes. For each cluster index $j = 1$ to $k - 1$:

1. Determine Target Size: Calculate the desired size for the current cluster, $s_j$: $s_j = \frac{|U_j|}{k-j+1}$ (This ensures the clusters are roughly equal in size.)

2. Select Seed: Choose a random starting geometric center (vertex) $v \in U_j$.
3. Grow community: Construct the cluster $C_j$ by starting a traversal at $v$ restricted to the subgraph induced by $U_j$. Perform a breadth-first search. Neighbors are visited in a random order. Stop adding nodes to $C_j$ once $|C_j| = s_j$ (or if the connected component runs out of nodes).
4. Update: Define the remaining set of nodes for the next iteration: $U_{j+1} = U_j / C_j$

After extracting steps 1 through $k-1$, assign all remaining nodes to the final cluster: $C_k = U_k$

##### 3.2 Binary activity matrix

To get the binary activity matrix $B \in \{0,1\}^{n \times m}$, we create a per-feature background signal vector $q$,

$$q = \lambda(Q_{20}(A_{\cdot j}))$$

where $Q_n$ is the nth quantile, and $\lambda$ a transformation function $\lambda(x) := \sqrt{1+x} - 1$
Then, the binary activity is given by

$$B_{ij} = A_{ij} > \alpha + \beta q_j$$

$$\therefore B = (A - \alpha) \cdot \text{diag}(\frac{1}{q}) - \beta > 0 \quad \{q > 0\}$$

The breadth of epigenetic marks is given by $\mathbf{1}^T B$.

#### 4 Metagene

To explore all the epigenetic patterns present in our data, we create metagenes.
We picked features based on one threshold per modality, and a second threshold on the summed modalities to capture all features where interesting patterns could exist in at least one modality. We created an embedding for features in the same way we did for the cells, that is, transposing the fragment matrix $X$, The only difference from creating the cell embedding is that we did not remove any PCs, since they did not correlate strongly with the total fragment counts of features.

#### 5 Inter-modality correlation

We were interested in how the different modalities we had access to correlated with each other. Since acetylation and methylation come from the same experiment and are paired, the correlation would be very easy to calculate. However, we were more interested in RNA expression patterns against our epigenetic marks. We availed ourselves of the joint landscape embedding we had already created.
A simple pearson or spearman correlation on our griddata wouldn't have given us the correlation score $\rho \in [-1, 1]^m$ that we wanted to create. This is because we wanted a score that would be sensitive to areas with high signal, and less sensitive to areas of the manifold where no cells had any signal. It would be easy to imagine a landscape where one cell type is highly expressed, and another cell type is highly methylated, but the correlation would be positive, because of all the other cell types that had a near-zero signal.
To make the correlation score, we first needed to normalize our griddata with respect to each gene.

##### 5.1 Griddata processing

Griddata $Z$ is created by process outlined in section 2.1. In addition to median and gaussian filter, we apply the following transformations: winsorization at quantile $q_\alpha$ to remove outliars at high ranges (can affect robustness of minmax), and minmax to range $[0, 1]$.

$$Z = \min(Z, Q_{q_\alpha}(Z))$$

$$Z = \frac{Z - \min(Z)}{\max(Z) - \min(Z)}$$

Then, we scaled all features to the same range, limiting the fold-upscale to a maximum of $\beta$

$$\nu_j = \frac{1}{Q_{q_\beta}(Z_{\cdot j})}$$

$$Z = Z \min(\nu, \beta)$$

Followed by a final winsorization $> q_f$ and minmax to range $[0, 1]$.

#### 5.2 Correlation score calculation

We create a weight $w \in [0,1]^{m \times g_x \times g_y}$ for our signal to calculate a weighted correlation between modalities. Since $Z \in [0,1]^{g_x \times g_y \times m}$, a multi-modal matrix can be described as $\mathcal{Z} \in [0,1]^{m \times 2 \times g_x \times g_y}$ where 2 is the number of modalities.

$$\rho_j = \frac{\text{Cov}(z_{j0}, z_{j1}; \text{weights} = w)}{\sigma_{z_{j0}} \sigma_{z_{j1}}}$$

where

$$w = \min(\sum_{\imath} z_{j\imath}^2, 1)^2$$

$$z_{j0} = \boldsymbol{vec}(\mathcal{Z}_{j0}), z_{j1} = \boldsymbol{vec}(\mathcal{Z}_{j1})$$

#### 5.3 RNA-epigenetic potential correlation

We wanted to calculate the correlation of epigenetic potential to RNA expression, And wanted it to be near -1 when the RNA was aligned with the minimum of the potential, and +1 when the RNA was aligned with the maximum of the potential, reflecting the potential itself, given by $f = \text{meth} - \text{acet}$. We chose to combine the correlations of acetylation to RNA, and methylation to RNA, since using the negative values given by $f$ would be impractical with our setup of weighted covariance. To do operations with the two correlations, we first used fisher z-transformation $z = \text{arctanh}(\rho)$, after which an inverse transformation would give us back a value in range of $[-1, 1]$,

$$\rho = \tanh(\frac{z_{meth} - z_{acet}}{2\tau})$$

Since the quality of the data didn't lead to acetylation or methylation $\rho$ being near -1 or 1, we added a coefficient $\tau = 2$ to scale the correlation.

### 6 Wild-type – knock-down normalization

We noticed that the KD mutant had different mean and variance of its per-cell-type fragment distribution. To compare fragments per cell type, we did z-normalization.

#### 6.1 Fragment count z-normalization

$$f_j = \log 2 \sum_{u \in C_j} \hat{X}_u$$

$$\hat{X}_{B_i} = X_{B_i} \frac{10^6}{\sum_{v \in B_i} X_v}$$

$$\hat{f}_j = \frac{(f_j - \mu_f)}{\sigma_f}$$

##### 6.2 Mean rank normalization

The KD mutant seemed also to have different enrichment of fragment count sorted by rank compared to wild-type.
Let $\mu_0, \mu_1 \in \mathbb{R}^d$ be the mean vectors for two distributions, and let $0 < \varepsilon \ll 1$ be a small threshold value. Define the placement permutation $\pi : \{1, \ldots, d\} \to \{1, \ldots, d\}$ such that:

$$\text{rank}(\mu_1(\pi(i))) = \text{rank}(\mu_0(i))$$

where ranks are assigned in ascending order. That is, $\pi(i)$ is the index in distribution 1 whose rank matches the rank of index $i$ in distribution 0.

$$\tilde{\mu}_0(i) = \max(\mu_0(i), \varepsilon), \quad \tilde{\mu}_1(i) = \max(\mu_1(i), \varepsilon)$$

Then, the diagonal matrix $D$ can be made,

$$D = \text{diag}(\text{clip}(\frac{\tilde{\mu}_0(i)}{\tilde{\mu}_1(\pi(i))}, s_{\text{min}}, s_{\text{max}}))$$

where $s_{min}$ and $s_{max}$ are the minimum and maximum scalar values.
Then, $X$ can simply be normalized by

$$\hat{X} = XD$$
